# Mettl3-mediated m^6^A modification of Fgf16 restricts cardiomyocyte proliferation during heart regeneration

**DOI:** 10.1101/2022.02.27.482166

**Authors:** Fu-Qing Jiang, Jia-Xuan Chen, Wu-Yun Chen, Wan-Ling Zhao, Guo-Hua Song, Chi-Qian Liang, Yi-Min Zhou, Huan-Lei Huang, Rui-Jin Huang, Hui Zhao, Kyu-Sang Park, Zhen-Yu Ju, Dong-Qing Cai, Xu-Feng Qi

## Abstract

Cardiovascular disease is the leading cause of death worldwide due to the inability of adult heart to regenerate after injury. *N*^6^-methyladenosine (m^6^A) methylation catalyzed by the enzyme methyltransferase-like 3 (Mettl3) plays important roles in various physiological and pathological bioprocesses. However, the role of m^6^A in heart regeneration remains largely unclear. To study m^6^A function in heart regeneration, we modulated Mettl3 expression in vitro and in vivo. Knockdown of *Mettl3* significantly increased the proliferation of cardiomyocytes and accelerated heart regeneration following heart injury in neonatal and adult mice. However, *Mettl3* overexpression decreased cardiomyocyte proliferation and suppressed heart regeneration in postnatal mice. Conjoint analysis of methylated RNA immunoprecipitation sequencing (MeRIP-seq) and RNA-seq identified *Fgf16* as a downstream target of Mettl3-mediated m^6^A modification during postnatal heart regeneration. RIP-qPCR and luciferase reporter assays revealed that Mettl3 negatively regulates *Fgf16* mRNA expression in an m^6^A-Ythdf2-dependent manner. The silencing of *Fgf16* suppressed the proliferation of cardiomyocytes. However, the overexpression of ΔFgf16, in which the m^6^A consensus sequence was mutated, significantly increased cardiomyocyte proliferation and accelerated heart regeneration in postnatal mice compared with wild-type Fgf16. Our data demonstrate that Mettl3 post-transcriptionally reduces *Fgf16* mRNA levels through an m^6^A-Ythdf2-dependen pathway, thereby controlling cardiomyocyte proliferation and heart regeneration.

## Introduction

Heart failure is a leading cause of morbidity and mortality worldwide, due to the inability to regenerate the injured heart after myocardial infarction (Narula *et al*. 1996; Bergmann *et al*. 2009; Bui *et al*. 2011). Although various strategies including cell-based and cell free therapies are being explored to promote heart regeneration in animal models and human patients (Lin & Pu 2014; Sahara *et al*. 2015; Arrell *et al*. 2020; Biressi *et al*. 2020), the efficacy of cardiac therapy and its clinical implications remain uncertain (Chong *et al*. 2014). It has been demonstrated that complete cardiac regeneration occurs in neonatal mouse heart after ventricular resection at postnatal day 1 (p1), but this capacity is lost at p7 (Porrello *et al*. 2011). It has been believed that cardiac regeneration in neonatal mice is primarily mediated by cardiomyocyte proliferation (Porrello *et al*. 2011; Mahmoud *et al*. 2013; Porrello *et al*. 2013). Therefore, it is crucial to explore the related signaling networks and underlying molecular mechanisms that are responsible for postnatal cardiomyocyte proliferation. Previously, many studies have focused on the signaling pathways contributing to the activation of proliferative factors that regulate gene expression in hearts (Mahmoud *et al*. 2013; Porrello *et al*. 2013; Feng *et al*. 2019). However, the potential roles of the post-transcriptional regulation of cardiac mRNAs in cardiac physiology and pathology are largely unknown. Previous studies have reported that mRNA abundance does not correlate with protein expression in failing hearts (Brundel *et al*. 2001; Su *et al*. 2015). Moreover, nuclear and cytosolic mRNA levels are not correlated in cardiomyocytes (Preissl *et al*. 2015). Therefore, these reports suggest that epitranscriptomic regulation plays an important role in healthy and pathological hearts.

Although more than 150 distinct chemical marks have been identified on cellular RNA (Boccaletto *et al*. 2018), the *N*^6^-methyladenosine (m^6^A) methylation catalyzed by the enzyme methyltransferase-like 3 (Mettl3) has been the most prevalent and best-characterized mRNA modification (Dominissini *et al*. 2012; Meyer *et al*. 2012; Fu *et al*. 2014). Recent studies have demonstrated that m^6^A plays an important role in gene regulation (Roundtree *et al*. 2017), animal development (Zhao *et al*. 2017; Frye *et al*. 2018), and human disease (Hsu *et al*. 2017; Chen *et al*. 2018; Paris *et al*. 2019). A number of studies have demonstrated that effectors of m^6^A pathways consist of “writers”, “erasers”, and “readers” that respectively install, remove, and recognize methylation (Shi *et al*. 2019). Growing evidence has revealed that m^6^A effectors in different biological systems are multifaced and tunable because their expression and cellular localization are cell type- and environmental stimuli-dependent (Dorn *et al*. 2019; Shi *et al*. 2019). Therefore, the mechanisms by which m^6^A regulate gene expression are complex and context-dependent.

Despite m^6^A mRNA methylation has been recognized as a posttranscriptional modification in multiple organisms including plants, yeast, flies, and mammals (Batista *et al*. 2014; Luo *et al*. 2014; Haussmann *et al*. 2016; Wei *et al*. 2018), the roles of this posttranscriptional process and functions of m^6^A mRNA methylation in cardiomyocyte proliferation and animal models of heart regeneration remain unknown. Here, we elucidate the landscape of m^6^A mRNA methylation during heart regeneration in neonatal mice and show that inhibition of the methylase Mettl3 promotes cardiomyocyte proliferation and heart regeneration. We further demonstrate that posttranscriptional downregulation of Fgf16 by Mettl3-mediated m^6^A modification via a YT521-B homology (YTH) domain family 2 (Ythdf2)-dependent pathway, inhibits heart regeneration in neonatal mice.

## Results

### Mettl3 expression and the m^6^A levels in mRNA are increased upon neonatal heart injury

It has been demonstrated that Mettl3-mediated m^6^A is enhanced in response to hypertrophic stimulation and is necessary for a normal cardiomyocyte hypertrophy in adult mice (Dorn *et al*. 2019), indicating the pivotal functions of Mettl3-mediated m^6^A in cardiac homeostasis. These further prompt us to ask whether m^6^A modification influences heart regeneration in neonatal mice. To determine the expression patterns of m^6^A methyltransferases (Mettl3 and Mettl14) and demethylase (Alkbh5 and Fto), myocardium isolated from neonatal hearts at a series of time points was subjected to quantitative real-time PCR (qPCR) assay. Among these four genes, Mettl3 expression significantly increased from postnatal day 3 (p3) to p14 (Figure 1A). In agreement with qPCR result, increased expression of Mettl3 in postnatal heart at p14 was also detected in protein levels (Figure 1, B and C). To determine the cardiac mRNAs modified by m^6^A methylation in neonatal mice, m^6^A levels were measured by an ELISA-based quantification assay at p3, p7, and p14. Increased levels of m^6^A modification were detected at p14 compared with p3 and p7 (Figure 1D), suggesting the lower levels of m^6^A modification within postnatal 7 days (the regenerative window). These results indicate the potential functional importance of the m^6^A modification of cardiac mRNAs for heart regeneration. To further determine the m^6^A modification in injured hearts, apical resection was performed at p3, followed by m^6^A level examination in sham and resected hearts at 5 days post-resection (dpr) (Figure 1E). As shown in Figure 1F, apex resection injury increased the level of m^6^A modification by 65% at 5 dpr compared with sham hearts. Moreover, the expression of *Mettl3* was approximately increased up to 2-fold at 5 dpr compared with sham hearts (Figure 1G). Western blotting revealed that the expression of Mettl3 rather than Mettl14 was greatly increased at 5 dpr compared with sham hearts (Figure 1, H-J). Taken together, these findings suggest that Mettl3-mediated m^6^A modification might be important for regulating heart regeneration.

**Figure 1.**
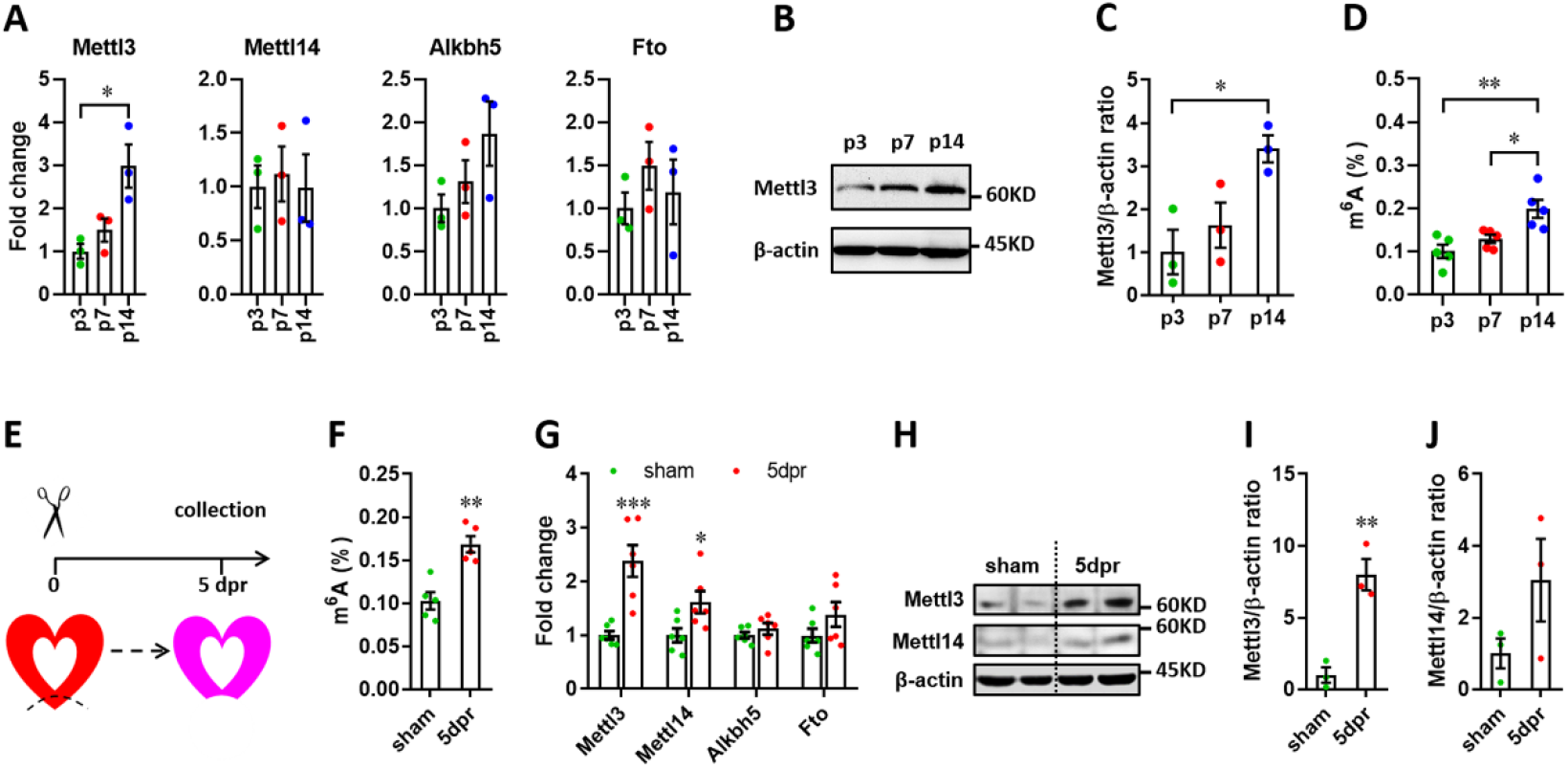
Expression patterns of m^6^A methylases and demethylases during heart regeneration in neonatal mice. (**A**) Quantification of m^6^A methylases (Mettl3 and Mettl14) and demethylases (Alkbh5 and Fto) expression in neonatal hearts at the indicated time points after birth (*n*=3 hearts). (**B** and **C**) Representative images (B) and quantification (C) of protein expression of Mettl3 in neonatal hearts (*n*=3 hearts). (**D**) Cardiac mRNA m^6^A levels in neonatal heart were measured by ELISA-based quantification assay (*n*=5 hearts per group). (**E**) Schematic of heart injury and sample collection in neonatal mice. (**F**) mRNA m^6^A levels in the injured hearts at 5 dpr were measured (*n*=5 hearts). (**G**) qPCR validation of m^6^A methylases and demethylases in neonatal hearts at 5 dpr (*n*=6 hearts). (**H-J**) Protein levels of Mettl3 and Mettl14 were measured by western blotting at 5 dpr. Representative images (F) and quantification (G and H) of protein expression of Mettl3 and Mettl14 are shown (*n*=3 hearts). All data are presented as the mean ± SEM. \**p*<0.05, \*\**p*<0.01, \*\*\**p*<0.001 compared with p3 (A and C) or sham (F, G and I). *P* values were determined by 1-way (A, C, and D) or 2-way (G) ANOVA with Dunnett’s multiple-comparison test, or by 2-tailed Student’s *t* test (F, I, and J).

### In vitro effects of Mettl3 on the proliferation of cardiomyocytes

Previous studies have demonstrated that cardiomyocyte proliferation is vital to hear regeneration (Porrello *et al*. 2011; Mahmoud *et al*. 2013). This idea promoted us to ask whether Mettl3-mediated m^6^A also influences cardiomyocyte proliferation. In the rat cardiomyocyte cell line (H9c2), we silenced the expression of *Metll3* gene using siRNA (siMettl3) (Supplemental Figure 1A) and performed nuclear incorporation assay with 5-ethynyl-2’-deoxyuridine (EdU). The percentage of EdU^+^ cells was significantly increased by *Mettl3* knockdown (Supplemental Figure 1, B and C), suggesting elevated levels of proliferation in *Mettl3*-silencing cells. Indeed, cell number was also greatly promoted by siMettl3 (Supplemental Figure 1D). To further explore the potential effects of Mettl3 overexpression on H9c2 cell proliferation, a stable cell line was established using lentiviral-mediated overexpression of Mettl3 (Supplemental Figure 1, E-H). As expected, increased m^6^A levels were observed in the Mettl3-overexpressing cells compared with controls (Supplemental Figure 1, I and J). In contrast with the knockdown cells, both cell proliferation and count were remarkably suppressed by Mettl3 overexpression in H9c2 cells (Supplemental Figure 1, K-M). These results reveal that cardiomyocyte proliferation might be negatively controlled by Mettl3-mediated m^6^A modification.

To further confirm this idea, primary cardiomyocytes isolated from neonatal mice were subjected to proliferation assay. Ki67 and cTnT double staining revealed that *Mettl3* knockdown (Figure 2, A and B) leads to an increase in the percentage of Ki67^+^ cTnT^+^ cells, indicating an elevated proliferation level of primary cardiomyocytes (Figure 2, E and F). In contrast, *Mettl3* overexpression (Figure 2, C and D) suppressed the proliferation of primary cardiomyocytes as demonstrated by Ki67 and cTnT double staining (Figure 2, G and H). In agreement with these results, EdU incorporation assay revealed that the percentage of EdU^+^ cTnT^+^ cells was greatly increased by *Mettl3* knockdown (Figure 2, I and J), indicating an elevated proliferation level of primary cardiomyocytes. However, the proliferation of primary cardiomyocytes was approximately suppressed by 48% by *Mettl3* overexpression as demonstrated by nuclear EdU incorporation (Figure 2, K and L). Taken together, these findings from primary cardiomyocytes and cell line suggest that Metll3-mediated m^6^A modification can negatively regulate the proliferation of cardiomyocytes.

**Figure 2.**
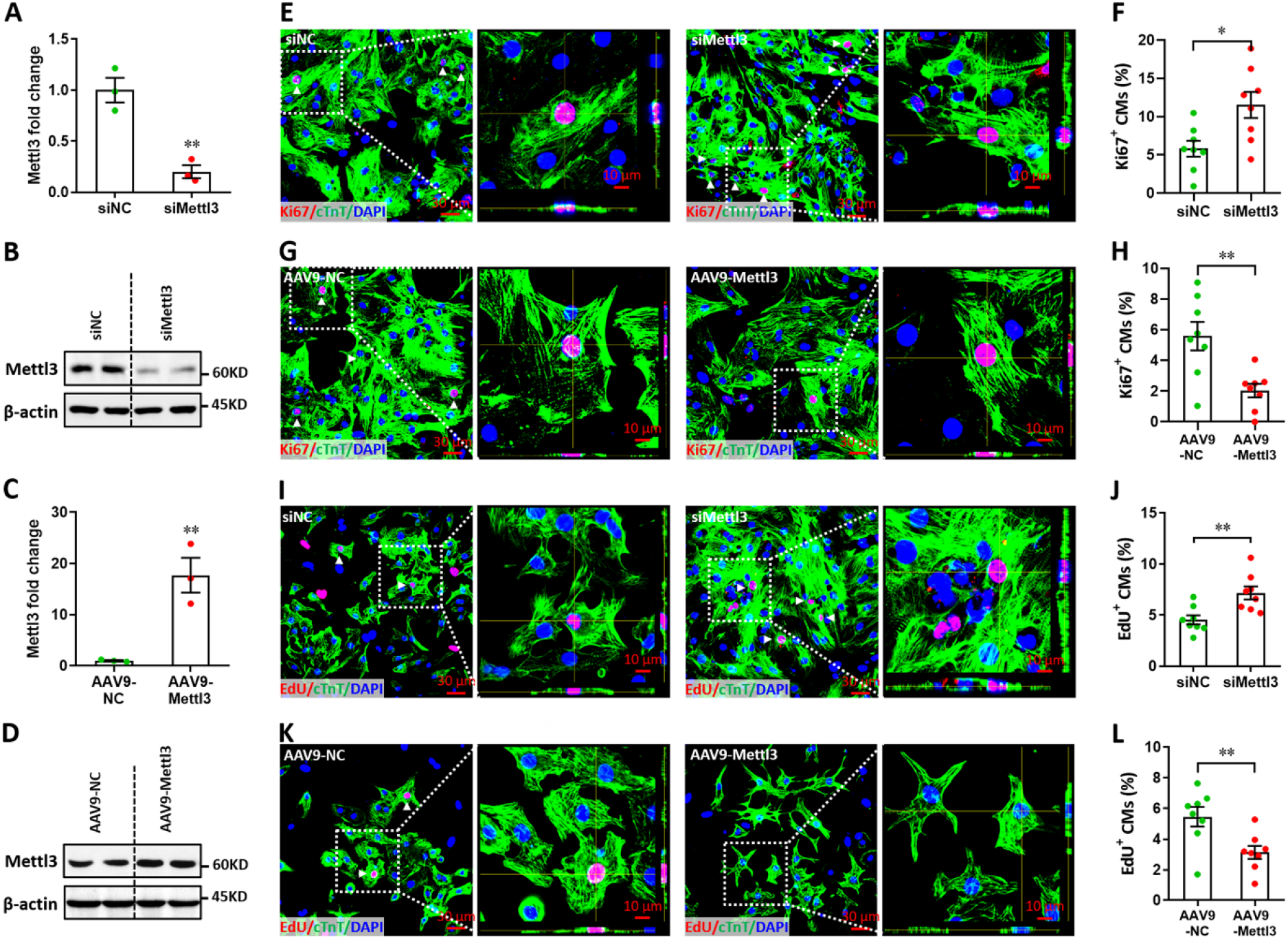
Mettl3 suppresses the proliferation of primary cardiomyocytes in vitro. (**A** and **B**) Primary cardiomyocytes were transfected with siMettl3 and siNC for 48 hr, followed by qPCR (A, *n*=3) and western blotting (B) validation. (**C** and **D**) Primary cardiomyocytes were transfected with AAV9-NC and AAV9-Mettl3 for 72 hr, followed by qPCR (C, *n*=3) and western blotting (D) validation. (**E-H**) Mettl3 silencing and overexpressing cardiomyocytes were subjected to Ki67 (red) and cTnT (green) double staining. Representative images (E, G) and quantification (F, H) of Ki67^+^ cardiomyocytes are shown (*n*=8). Magnified Z-stack confocal images of Ki67^+^ cardiomyocytes are shown in right panel (E, G). (**I-L**) Mettl3 silencing and overexpressing cardiomyocytes were subjected to EdU (red) and cTnT (green) double staining. Representative images (I, K) and quantification (J, L) of EdU^+^ cardiomyocytes are shown (*n*=8). Magnified Z-stack confocal images of EdU^+^ cardiomyocytes are shown in right panel (I, K). All data are presented as the mean ± SEM of three separate experiments, \**p*<0.05, \*\**p*<0.01 versus control. *P* values were determined by 2-tailed Student’s *t* test.

### In vivo knockdown of Mettl3 promotes cardiomyocyte proliferation in injured neonatal hearts

To knock down Mettl3 in myocardium of neonatal mice, adeno-associated virus 9 (AAV9)-mediated delivery of shRNA was used. Firstly, the time course efficiency of AAV9-shRNA system was determined in neonatal heart using the harboring EGFP reporter gene. Our results revealed that AAV9-shRNA system induces modest expression of reporter gene in neonatal heart at 2 days post-injection (dpi), followed by substantial expression from 6 to 27 dpi (Supplemental Figure 2). To investigate the *in vivo* roles of Mettl3 in cardiomyocyte proliferation, AAV9-shMettl3 viruses were injected into neonatal hearts at p1, followed by apex resection at p3 (equal to 0 dpr) and a nuclear EdU incorporation assay at 5 dpr (Figure 3A). Successful knockdown of Mettl3 in heart tissue at 0 dpr was confirmed by the qPCR validation of Mettl3 mRNA expression (Figure 3B). To further confirm Mettl3 knockdown, primary cardiomyocytes isolated from ventricles at 0 dpr were subjected to western blotting and ELISA-based m^6^A quantification assay. The decreased expression of Mettl3 protein (Figure 3, C and D) and reduced m^6^A modification (Figure 3E) were further demonstrated in primary cardiomyocytes. To determine the size of the cardiomyocytes in neonatal heart at 5 dpr, wheat germ agglutinin (WGA) staining was performed. The cardiomyocyte size was smaller in Mettl3-silencing hearts compared with controls (Figure 3, F and G), implying increased numbers of cardiomyocytes in Mettl3-deficient hearts. These results suggest that *in vivo* deficiency of Mettl3 promotes heart regeneration, implying elevated proliferation potential of cardiomyocytes.

**Figure 3.**
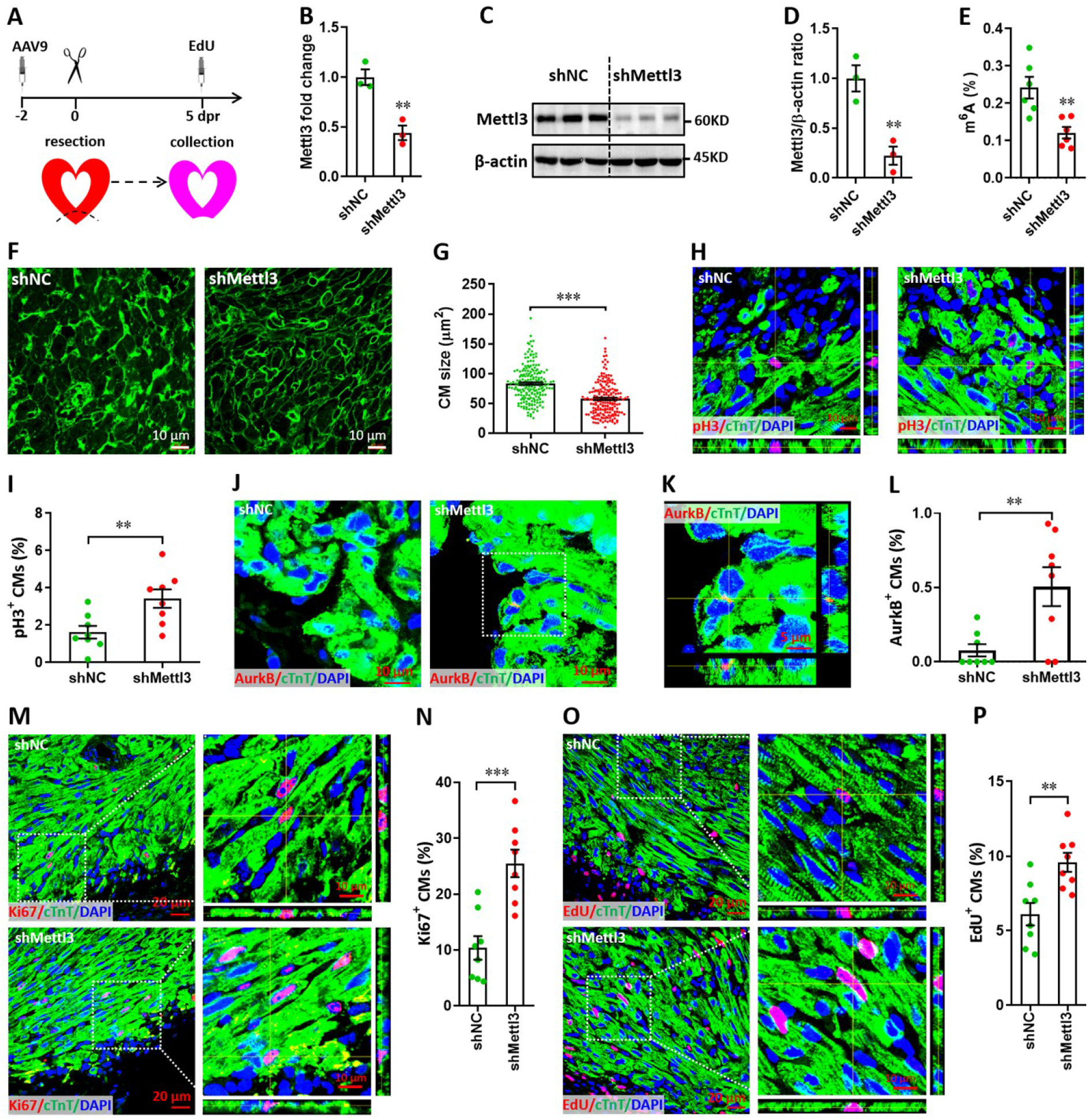
Mettl3 knockdown promotes cardiomyocyte proliferation in neonatal hearts at 5 dpr. (**A**) Schematic of AAV9-shMettl3 virus injection designed to knock down *Mettl3* in neonatal hearts. (**B**) qPCR validation of *Mettl3* in the AAV9-injected hearts at 0 dpr (*n*=3 hearts). (**C-E**) Primary cardiomyocytes were isolated from neonatal hearts with AAV9 virus injection at 0 dpr, followed by western blotting and m^6^A quantification. Representative images (C) and quantification (D, *n*=3) of Mettl3 protein expression, as well as the quantification of m^6^A modification levels (E, *n*=6) are shown. (**F** and **G**) Representative WGA staining images (F) and quantification (G) of the size of cardiomyocytes located in border zone at 5 dpr (*n*=∼200 cells from 5 hearts per group). (**H** and **I**) Representative Z-stack confocal images (H) and quantification (I) of pH3^+^ cardiomyocytes in apical ventricle at 5 dpr (*n*=8 hearts). (**J**-**L**) Representative images (J and K) and quantification (L) of AurkB^+^ cardiomyocytes in apical zone at 5 dpr (*n*=8 hearts). K, magnified Z-stack confocal image in J (right panel). (**M** and **N**) Representative images (M) and quantification (N) of Ki67^+^ cardiomyocytes in apical ventricle at 5 dpr (*n*=8 hearts). Right panel (M), magnified Z-stack confocal images. (**O** and **P**) Representative images (O) and quantification (P) of EdU^+^ cardiomyocytes in apical ventricle at 5 dpr (*n*=8 hearts). Right panel (O), magnified Z-stack confocal images. All data are presented as the mean ± SEM, \**p*<0.05, \*\**p*<0.01, \*\*\**p*<0.001 versus control. *P* values were determined by 2-tailed Student’s *t* test.

To further confirm this conjecture, in vivo proliferation of cardiomyocytes was further examined at 5 dpr. It has been demonstrated that cardiomyocytes can go through cell cycle phases but stop and not complete it (Lazar *et al*. 2017). To prove complete cardiomyocyte proliferation, late cell cycle markers including phospho-Histone H3 (pH3) and aurora kinase B (AurkB) were further used in this study. Consistently, the percentage of pH3^+^ cTnT^+^ cells was significantly elevated by Mettl3 silencing both in apical (Figure 3, H and I) and remote (Supplemental Figure 3, A and B) zones, indicating that Mettl3 knockdown really promotes the complete proliferation of cardiomyocytes *in vivo*. Importantly, increased percentage of AurkB^+^ cardiomyocytes was also detected in cardiac apex in the Mettl3-deficient neonatal mice (Figure 3, J-L), implying the increased cytokinesis of cardiomyocytes. In agreement with above observation, in vivo staining of Ki67 further revealed that cardiomyocyte proliferation in the injured neonatal hearts was greatly increased by Mettl3 silencing in the apical (Figure 3, M and N) and remote (Supplemental Figure 3, C and D) zones compared with controls. Moreover, nuclear EdU incorporation revealed that *in vivo* silencing of *Mettl3* increased the percentage of EdU^+^ cTnT^+^ cells both in apical (Figure 3, O and P) and remote (Supplemental Figure 3, E and F) cardiac tissues. These data suggest that Mettl3-mediated m^6^A modification might negatively controls cardiomyocyte proliferation during neonatal heart regeneration. Taken together, these in vitro and in vivo findings strongly demonstrated that Mettl3-mediated m^6^A modification can negatively regulate cardiomyocyte proliferation during heart regeneration in neonatal mice.

To further explore whether Mettl3 knockdown influences cardiomyocyte proliferation in the homeostatic neonatal hearts without injury, AAV9-shMettl3 and control viruses were injected at p1, followed by sample collection and histological analysis at p8 (Supplemental Figure 4A). This virus injection strategy and expression time are equal to the time point of 5 dpr in the apex resection injury model as mentioned in Figure 3A. Our data showed that Mettl3 knockdown has no significant effects on heart weight/body weight (HW/BW) ratio compared with controls (Supplemental Figure 4B). WGA staining revealed that cardiomyocyte size is comparable between Mettl3-silencing and control hearts (Supplemental Figure 4, C and D). Moreover, no significant differences in cardiomyocyte proliferation were observed in Mettl3-silencing hearts compared with controls, which were determined by EdU, Ki67, and pH3 staining (Supplemental Figure 4, E-J). These findings suggest that Mettl3 knockdown has no significant effects on cardiomyocyte proliferation in the uninjured neonatal hearts.

### Mettl3 knockdown accelerates heart regeneration in neonatal mice

It is well known that neonatal mice possess the capacity of heart regeneration upon injury within p7. However, neonatal mouse could not completely finish the heart regeneration until 21 days post-injury even induced at p1 (Porrello *et al*. 2011; Mahmoud *et al*. 2013; Porrello *et al*. 2013). To further investigate whether Mettl3 deficiency indeed promotes heart regeneration, AAV9-shMettl3 viruses were injected into neonatal mice at p1, followed by apex resection at p3 and sample collection at 14 dpr, a time point prior to the completion of heart regeneration (Figure 4A). Histological analysis revealed that scar size in the cardiac apex at 14 dpr was decreased by 73% in Mettl3-deficient hearts compared with controls (Figure 4, B and C), implying the accelerated heart regeneration. In agreement with these results, smaller scar sizes in Mettl3-deficient hearts were also confirmed by whole heart images (Figure 4, D-F). In addition, cardiomyocyte size was much smaller in Mettl3-deficient hearts compared with controls (Figure 4, G and H). These results suggest that Mettl3 deficiency leads to fast regeneration of neonatal heart as early as 14 dpr. As expected, increased left ventricular ejection fraction (LVEF) and fractional shortening (LVFS) were detected in Mettl3-deficient mice (Figure 4, I-K), indicating that left ventricular systolic function was promoted by Mettl3 knockdown. To determine proliferating cardiomyocytes during the whole period of heart regeneration within 14 dpr, EdU was injected in a pulse chase manner (Figure 4A). Our data revealed that the percentage of EdU^+^ cTnT^+^ cells both in apical and remote zones were significantly increased in Mettl3-deficient hearts compared with controls (Figure 4, L-O), implying that there are more proliferating cardiomyocytes during the whole period of 14 dpr. Taken together, these data suggest that Mettl3 knockdown promotes cardiomyocyte proliferation in the early stage of heart regeneration, thereby accelerating heart regeneration within two weeks post-resection in neonatal mice. In contract, Mettl3 knockdown did not lead to significant changes in cardiomyocyte size, proliferation, as well as cardiac function in the homeostatic neonatal mice without injury (Supplemental Figure 5).

**Figure 4.**
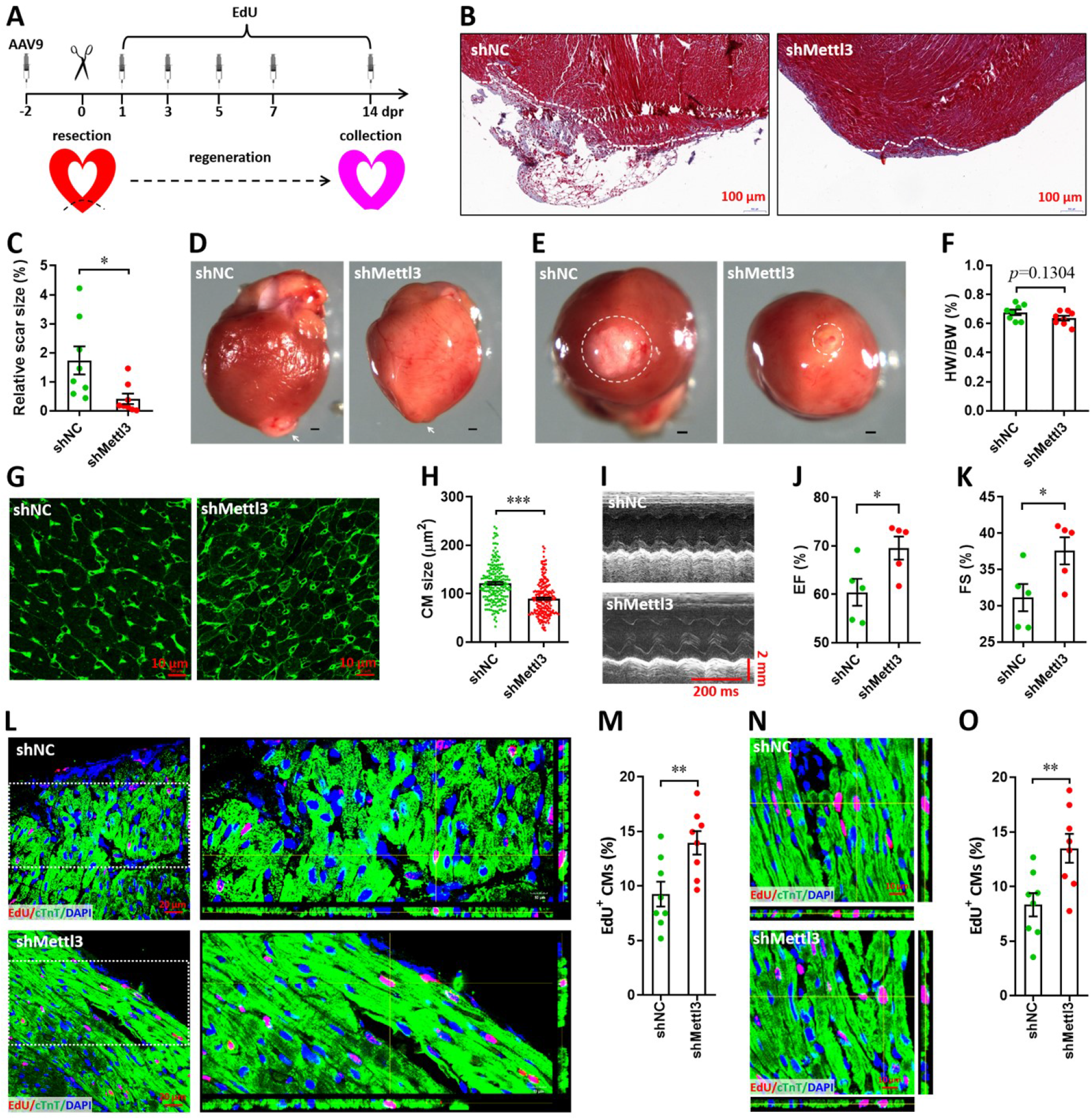
Knockdown of Mettl3 accelerates heart regeneration in neonatal mice at 14 dpr. (**A**) Schematic of AAV9 virus injection, apex resection, and EdU-pulse injection, followed by sample collection at 14 dpr. (**B** and **C**) Representative masson’s trichrome staining images of cardiac apex (B) and quantification of scar size (C) in control and Mettl3-deficient hearts (*n*=8 hearts). (**D** and **E**) Representative whole (D) and apical (E) images of neonatal hearts at 14 dpr. Arrowhead (D) and circle (E) denote scars in cardiac apex. (**F**) Quantification of heart weight (HW) to body weight (BW) ratio (*n*=8 hearts). (**G**) Representative WGA staining images of myocardium located in border zone in control and Mettl3-deficient hearts. (**H**) Quantification of cardiomyocyte size in control and Mettl3 knockdown hearts are shown (*n*=∼200 cells from 5 hearts). (**I**-**K**) Representative images of M-model echocardiography (I) and quantification of LVEF (J) and LVFS (K) are shown (*n*=6 hearts). (**L**) Representative images of EdU^+^ cardiomyocytes (double positive for EdU and cTnT) in the injured cardiac apex. Right panel, representative Z-stack confocal images. (**M**) Quantification of EdU^+^ cardiomyocytes in apical zone of control and Mettl3-deficient hearts (*n*=8 hearts). (**N** and **O**) Representative Z-stack confocal images (N) and quantification (O) of EdU^+^ cardiomyocytes in the uninjured ventricle (remote zone) are shown (*n*=8 hearts). All data are presented as the mean ± SEM, \**p*<0.05, \*\**p*<0.01, \*\*\**p*<0.001 versus control. *P* values were determined by 2-tailed Student’s *t* test.

To further explore whether Mettl3 knockdown extends the regenerative window, AAV9-shMettl3 viruses were injected at p1, followed by apex resection at p7, when the regenerative potential has mostly ceased (Supplemental Figure 6A). Mettl3 knockdown at p7 was confirmed by western blotting (Supplemental Figure 6, B and C). Moreover, Mettl3 knockdown-induced decrease in m^6^A levels at p7 was further demonstrated by ELISA-based m^6^A quantification assay (Supplemental Figure 6D). To determine the proliferation of cardiomyocytes at 3 dpr (the early stage of heart injury), immunofluorescence staining was performed using antibodies against Ki67 or pH3 together with cTnT antibodies. Ki67 staining revealed that cardiomyocyte proliferation in the apical zones of injured neonatal hearts was significantly increased by Mettl3 knockdown (Supplemental Figure 6, E-G). In addition, Mettl3-silencing increased the percentage of pH3^+^ cTnT^+^ cells in apical myocardium (Supplemental Figure 6, H-J). These results are agreement with the histological analysis at 21 dpr which revealed that scar size in the cardiac apex was decreased in Mettl3-deficient hearts compared with controls (Supplemental Figure 6K), implying the improved heart regeneration. Moreover, improving cardiac function was detected in Mettl3-deficient heart at 21 dpr as demonstrated by the increased LVEF and LVFS values (Supplemental Figure 6, L-N). Taken together, these findings further suggest that Mettl3 deficiency extends regenerative window in neonatal mice.

### Mettl3 overexpression suppresses heart regeneration in postnatal mice

We subsequently investigated the potential roles of Mettl3 overexpression in heart regeneration. To overexpress Mettl3 in cardiomyocytes, AAV9-Mettl3 viruses were injected into neonatal mice at p1, followed by apex resection at p3 (Figure 5A). Overexpression of Mettl3 protein and increase in m^6^A modification at p3 was confirmed by western blotting (Figure 5, B and C) and ELISA-based m^6^A quantification assay (Figure 5D), respectively. To determine the potential inhibition of heart regeneration induced by Mettl3 overexpression, the resected neonatal mice were raised until 28 dpr (Figure 5A), when beyond the time to complete regeneration (21 dpr) for neonatal mice (Porrello *et al*. 2011). Persistent overexpression of Mettl3 at 28 dpr was further evidenced by qPCR and immunofluorescent staining (Supplemental Figure 7, A and B), this was further supported by the substantial expression of EGFP reporter gene (Supplemental Figure 7C). Trichrome staining results revealed an increase in fibrotic scar size in Mettl3-overexpressing hearts compared with controls (Figure 5, E-G). The larger scar size induced by Mettl3 overexpression was further confirmed by whole heart images in parallel with the increased HW/BW ratio (Figure 5, H-J). In agreement with these results, cardiomyocyte size was increased in response to Mettl3 overexpression at 28 dpr (Figure 5, K and L). To evaluate the effects of Mettl3 on cardiac function, mice were subjected to echocardiography. Our data showed that Mettl3 overexpression leads to decreases in LVEF and LVFS levels (Figure 5, M-O), indicating that left ventricular systolic function is attenuated by Mettl3 overexpression.

**Figure 5.**
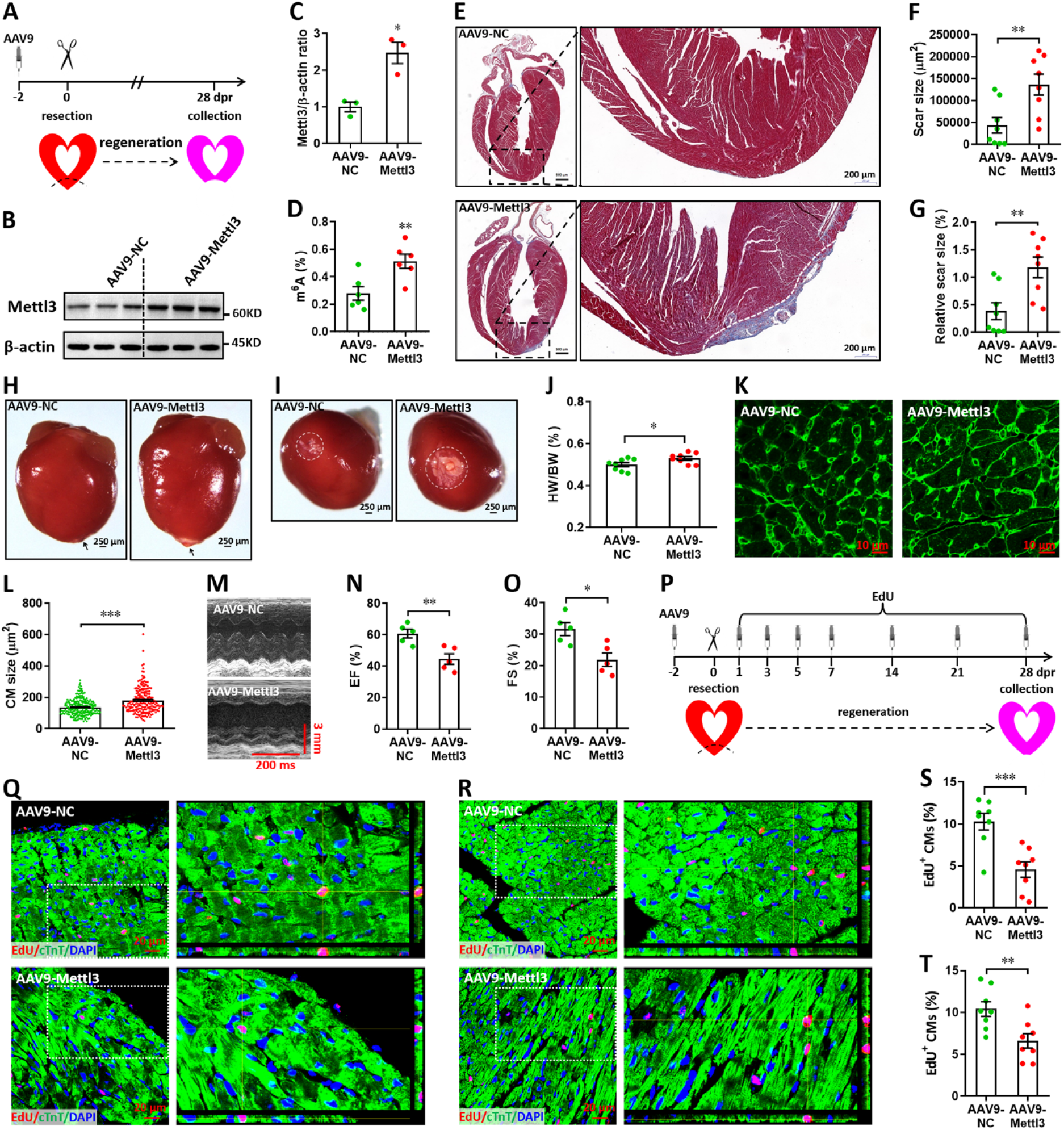
Overexpression of Mettl3 suppresses heart regeneration in neonatal mice upon injury. (**A**) Schematic of AAV9-Mettl3 virus injection, apex resection, and sample collection in neonatal mice. (**B** and **C**) Representative images (B) and quantification (C) of western blotting for Mettl3 expression in neonatal heart at 0 dpr (*n*=3 hearts). (**D**) m^6^A levels in the heart at 0 dpr were measured by ELISA-based quantification assay (*n*=6 hearts). (**E**) Representative masson’s trichrome staining images of heart in control and Mettl3-overexpressing mice at 28 dpr. (**F** and **G**) Direct (F) and relative (G) quantification of scar size in control and Mettl3-overexpressing hearts (*n*=8 hearts). Relative scar size is presented as percentages of whole ventricle size. (**H** and **I**) Representative whole (H) and apical (I) images of neonatal hearts at 28 dpr. Arrowhead (H) and circle (I) denote scars in cardiac apex. (**J**) Quantification of heart weight (HW) to body weight (BW) ratio (*n*=8 hearts). (**K** and **L**) Representative WGA staining images (K) and quantification of cardiomyocyte size (L) in border zone of control and Mettl3-overexpressing hearts at 28 dpr (*n*=∼200 cells from 5 hearts). (**M**-**O**) Representative images of M-model echocardiography (M) and quantification of LVEF (N) and LVFS (O) are shown at 28 dpr (*n*=8 hearts). (**P**) Schematic of EdU pulse injection and sample collection at 28 dpr. (**Q** and **R**) Representative images of EdU^+^ cardiomyocytes in apical (Q) and remote (R) zone at 28 dpr. Right panel, magnified Z-stack confocal images. (**S** and **T**) Quantification of EdU^+^ cardiomyocytes in apical (S) and remote (T) zone in control and Mettl3-overexpressing hearts (*n*=8 hearts). All data are presented as the mean ± SEM, \**p*<0.05, \*\**p*<0.01, \*\*\**p*<0.001 versus control. *P* values were determined by 2-tailed Student’s *t* test.

Using the immunofluorescence staining with antibodies against Ki67 and pH3, no positive cardiomyocytes were detected in both groups at 28 dpr (Supplemental Figure 8). However, pulse-chase injection of EdU (Figure 5P) revealed that cardiomyocyte proliferation in the apical (Figure 5, Q and S) and remote (Figure 5, R and T) zones was remarkably suppressed by Mettl3 overexpression during the whole period of 28 days post-resection. Therefore, these results suggest that Mettl3 overexpression might suppress heart regeneration in neonatal mice by inhibiting cardiomyocyte proliferation in the early stage of hear injury and repair. However, Mettl3 overexpression did not change the HW/BW ratio and cardiomyocyte size in the postnatal mice without injury (Supplemental Figure 9, A-D). EdU-pulse injection experiments showed that cardiomyocyte proliferation was not influenced by Mettl3 overexpression in the uninjured hearts (Supplemental Figure 9, E and F). In line with these results, there were no significant changes in cardiac function in the uninjured hearts upon Mettl3 overexpression (Supplemental Figure 9, G and H).

### Mettl3 knockdown promotes heart regeneration at non-regenerative stages

Subsequently, myocardium infarction (MI) injury model was used in postnatal and adult mice to further determine the involvement Mettl3 in heart regeneration at the non-regenerative stages. To silence Mettl3 expression in heart, AAV9-shMettl3 and control viruses were injected into neonatal mice at p1, followed by MI surgery at p7. Samples were then collected and analyzed at 5 and 21 dpM, respectively (Figure 6A). WGA staining results revealed that Mettl3 deficiency significantly reduced the size of cardiomyocytes in the infarcted neonatal hearts at 5 dpM (Figure 6, B and C). Ki67 and cTnT double staining revealed that Mettl3 knockdown promotes the proliferation of cardiomyocytes in the border zone of infarcted neonatal hearts as evidenced by the increased percentage of Ki67^+^ cTnT^+^ cells (Figure 6, D and E). Moreover, increased percentage of pH3^+^ cTnT^+^ cells was detected in the border zone in the Mettl3-deficient hearts at 5 dpM (Figure 6, F and G). We further analyzed the cardiac histology at 21 dpM and found that both HW/BW ratio and scar size were lower in the Mettl3-deficient heart compared with controls (Figure 6, H-J). In agreement with these findings, smaller size of cardiomyocytes was also detected in the Mettl3-deficient hearts compared with controls (Figure 6, K and L). These results suggest that Mettl3 deficiency might attenuate the cardiac remodeling during myocardium infarction in postnatal mice. Echocardiography analysis at 21 dpM showed that Mettl3 deficiency significantly increases LVEF and LVFS levels compared with control hearts (Figure 6, M-O).

**Figure 6.**
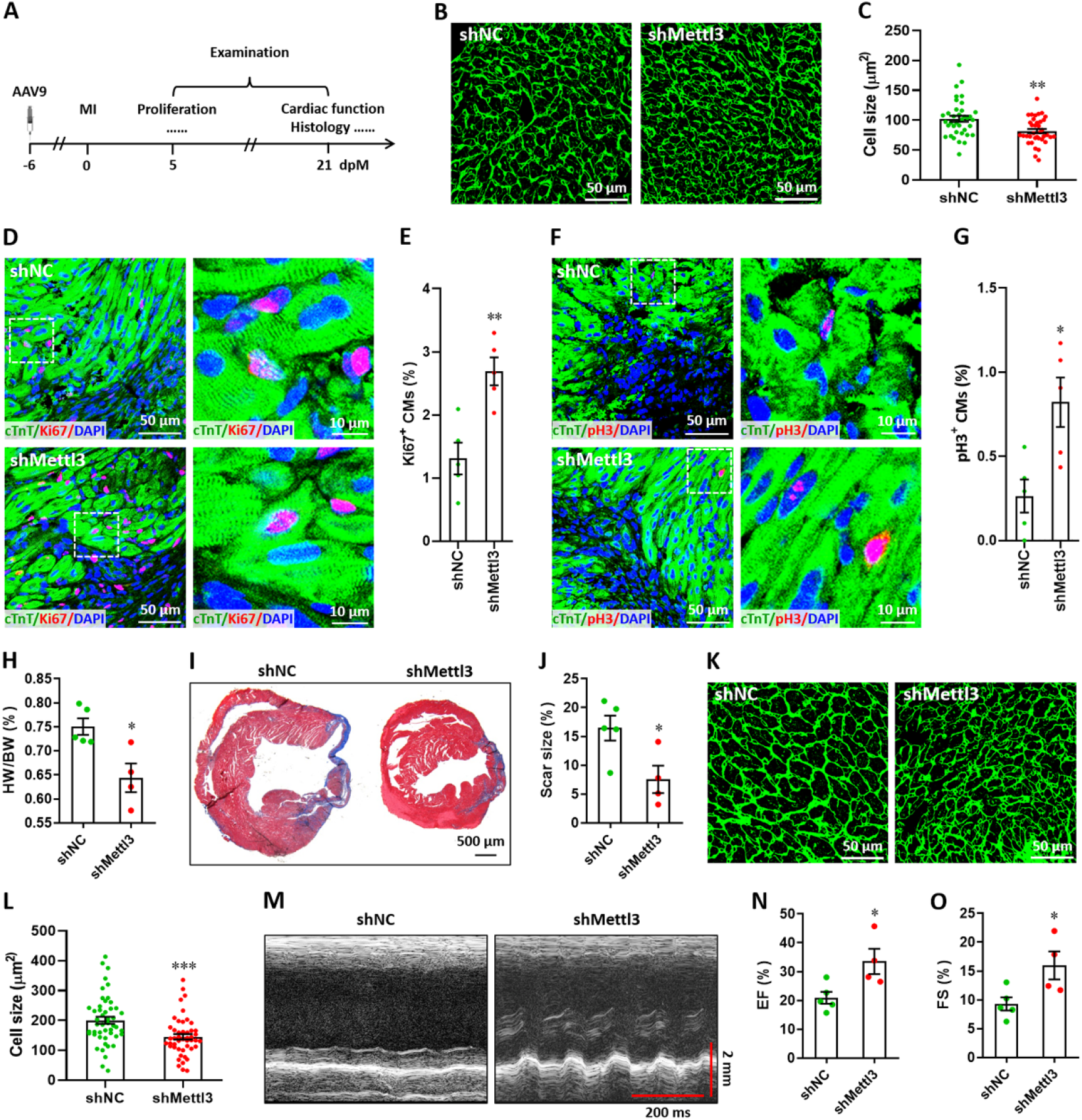
Mettl3 deficiency promotes heart regeneration in postnatal mice injured at p7. (**A**) Schematic of AAV9 virus injection at p1, myocardium infarction (MI) induction at p7, and histological analysis at 5 and 21 dpM. (**B** and **C**) Representative WGA staining images (B) and quantification of cardiomyocyte size (C) in control and Mettl3-deficient hearts at 5 dpM (*n*=∼100 cells from 5 hearts). (**D** and **E**) Representative images (D) and quantification (E) of Ki67^+^ cardiomyocytes in the border zone of injured hearts at 5 dpM (*n*=5 hearts). (**F** and **G**) Representative images (F) and quantification (G) of pH3^+^ cardiomyocytes in the border zone of injured hearts at 5 dpM (*n*=5 hearts). (**H**) Quantification of heart weight (HW) to body weight (BW) ratio at 21 dpM (*n*=5 hearts). (**I** and **J**) Representative masson’s trichrome staining images (I) and quantification of scar size (J) in control and Mettl3-deficient hearts at 21 dpM (*n*=5 hearts). (**K** and **L**) Representative WGA staining images (K) and quantification of cardiomyocyte size (L) in control and Mettl3-deficient hearts at 21 dpM (*n*=∼100 cells from 5 hearts). (**M**-**O**) Representative images of M-model echocardiography (M) and quantification of LVEF (N) and LVFS (O) are shown at 21 dpM (*n*=5 hearts). All data are presented as the mean ± SEM, \**p*<0.05, \*\**p*<0.01, \*\*\**p*<0.001 versus control. *P* values were determined by 2-tailed Student’s *t* test.

In addition, we further confirmed our conclusion in adult mice model. To knock down the expression of Mettl3 in the heart of adult mice, 8-week-old mice were injected with AAV9-shMettl3 and control viruses, respectively. Mice were then subjected to MI surgery 4 weeks after virus injection, followed by histological analysis and cardiac function analysis at 14 and 28 dpM, respectively (Figure 7A). The persistent expression of AAV9-shMettl3 in adult heart after virus injection for 4 weeks (equal to 0 dpM) was validated by the immunofluorescent staining for EGFP (Figure 7B). At 14 dpM, HW/BW ratio and scar size were decreased by Mettl3 deficiency compared with controls (Figure 7, C-E). In line with these results, the decreased HW/BW ratio and scar size were also observed in the Mettl3-deficient hearts compared with controls at 28 dpM (Figure 7, F-H). These results suggest that Mettl3 deficiency might attenuate the cardiac remodeling during myocardium infarction. To further evaluate the effects of Mettl3 deficiency on cardiac function, mice were subjected to echocardiography analysis at 0 and 28 dpM, respectively. Our data showed that Mettl3 deficiency leads to a little increase in LVEF and LVFS levels at 0 dpM (prior to MI). However, increased levels of LVEF and LVFS were detected in Mettl3-deficient hearts at 28 dpM compared with control hearts, respectively (Figure 7, I-K). These results indicate that Mettl3 deficiency can significantly attenuate the cardiac dysfunction induced by MI in adult mice, but this benefit is not sensitive in homeostatic hearts.

**Figure 7.**
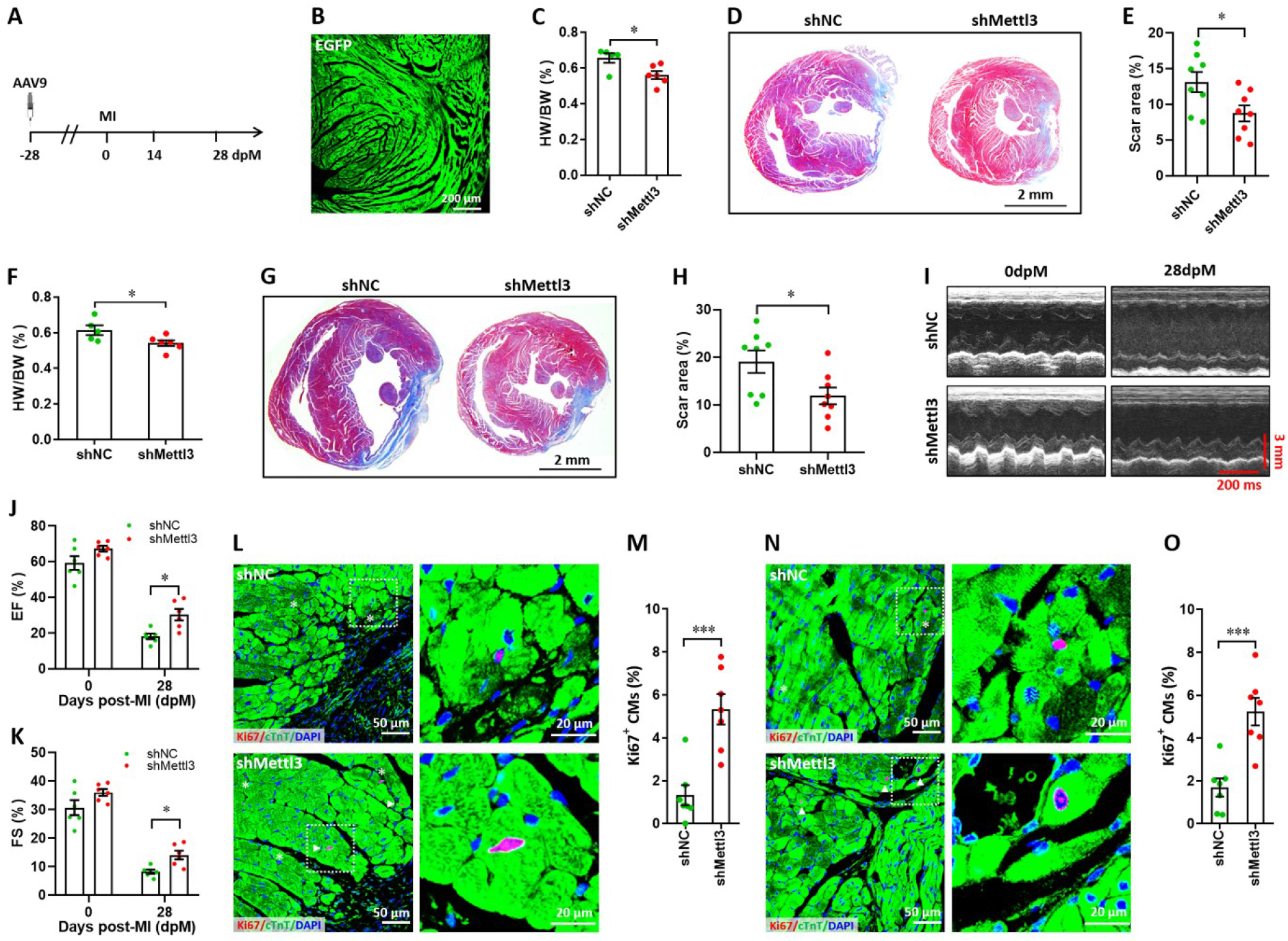
Mettl3 deficiency promotes heart regeneration in adult mice upon myocardium infarction injury. (**A**) Schematic of AAV9 virus injection 4 weeks prior to myocardium infarction (MI), followed by histological analysis at 14 and 28 dpM. (**B**) Representative images of reporter gene (EGFP) expression mediated by AAV9-shMettl3 virus in heart at 0 dpM. (**C**) Quantification of heart weight (HW) to body weight (BW) ratio at 14 dpM (*n*=6 hearts). (**D** and **E**) Representative masson’s trichrome staining images (D) and quantification of scar size (E) in control and Mettl3-deficient hearts at 14 dpM (*n*=8 hearts). (**F**) Quantification of HW to BW ratio at 28 dpM (*n*=6 hearts). (**G** and **H**) Representative masson’s trichrome staining images (G) and quantification of scar size (H) in control and Mettl3-deficient hearts at 28 dpM (*n*=8 hearts). (**I**-**K**) Representative images of M-model echocardiography (I) and quantification of LVEF (J) and LVFS (K) are shown at 0 and 28 dpM (*n*=6 hearts). (**L** and **M**) Representative images (L) and quantification (M) of Ki67^+^ cardiomyocytes in the border zone of injured hearts at 14 dpM (*n*=7 hearts). (**N** and **O**) Representative images (N) and quantification (O) of Ki67^+^ cardiomyocytes in the remote zone of injured hearts at 14 dpM (*n*=7 hearts). All data are presented as the mean ± SEM, \**p*<0.05, \*\*\**p*<0.001. *P* values were determined by 2-tailed Student’s *t* test (C, E, F, H, M, and O), or by 2-way ANOVA with Dunnett’s multiple-comparison test (J and K).

To further determine whether Mettl3 deficiency influences the proliferation of cardiomyocytes in adult mice during MI injury, immunofluorescent staining was performed in cardiac tissues at 14 dpM. Ki67 staining results showed that Mettl3 deficiency greatly increased the proliferation of cardiomyocytes in the border zone of infarction, as indicated by the increased percentage of Ki67^+^ cTnT^+^ cells (Figure 7, L and M). In consistent with border zone, increased percentage of Ki67^+^ cTnT^+^ cells was also detected in the Mettl3-deficient hearts in the remote zone in comparison with control hearts (Figure 7, N and O). These above findings imply that Mettl3 deficiency promotes cardiac repair and regeneration in postnatal mice at the non-regenerative stages, at least in part, through promoting cardiomyocyte proliferation.

### Variations in m^6^A-regulated heart genes upon injury

To investigate target genes involved in both Mettl3-mediated m^6^A modification and heart regeneration, ventricles from sham and 5 dpr hearts were subjected to methylated RNA immunoprecipitation sequencing (MeRIP-seq) and RNA-seq analysis. MeRIP-seq revealed that the GGACU motif is highly enriched in m^6^A sites in both sham and apex-resected hearts (Supplemental Figure 10A). Moreover, m^6^A peaks are particularly abundant in the vicinity of stop codons (Supplemental Figure 10B). However, the m^6^A peak distribution decreased in 5’ untranslated region (5’UTR) in apex-resected hearts, whereas there was an increase in coding regions (CDS) and stop codon regions compared with sham operated hearts (Supplemental Figure 10B). We further investigated the m^6^A distribution patterns in total and unique peaks. In contrast to total peaks which had an identical distribution, unique injury-dependent peaks demonstrated a distinct pattern in which a great increase in m^6^A deposits appeared in the CDS together with a relative decrease in both 5’UTRs and non-coding RNAs (Supplemental Figure 10C). As expected, an increase in total m^6^A peaks and m^6^A-tagged mRNAs was detected in injured hearts (Supplemental Figure 10D). In a comparison of the sham-operated and apex-resected hearts, a total of 750 genes (635 upregulated and 115 downregulated) were identified with a 2-fold m^6^A change (Supplemental Figure 10E). Gene Ontology (GO) analysis of these 635 upregulated genes indicated that a handful of the genes were associated with cardiac formation and assembly, hippo signaling, and establishment of spindle orientation (Supplemental Figure 10F).

In addition, both m^6^A-tagged transcripts overlapped between two different repeats and total MeRIP-seq reads were increased in the resected hearts compared with sham (Supplemental Figure 11, A and B), indicating an increase in m^6^A-tagged transcripts in injured hearts. A substantial proportion (about 60%) of total m^6^A-tagged transcripts (1,458 of 2,511) were specifically detected in the resected rather than sham hearts (Supplemental Figure 11C), implying that heart injury results in different landscapes of m^6^A modification. Compared with the 1,053 overlapping transcripts, GO analysis revealed that there were different GO biological terms including cell cycle for the 1,458 transcripts specifically tagged by m^6^A in injured hearts (Supplemental Figure 11, D and E).

### Mettl3-mediated m^6^A modification downregulates Fgf16 expression in cardiomyocytes

RNA-seq analysis identified 3,213 upregulated and 809 downregulated genes at 5 dpr compared with sham group (Supplemental Figure 12, A and B). Gene-set enrichment analysis (GSEA) revealed that cardiac muscle cell development and proliferation gene sets are related to neonatal heart injury and regeneration (Supplemental Figure 12, C and D). Based on RNA-seq and m^6^A-seq data, 333 overlapped genes between the m^6^A-tagged and mRNA-upregulated genes were identified at 5 dpr (Supplemental Figure 13A). GO analysis revealed that chromatin structure adjustment and cell cycle are important for cardiac regeneration (Supplemental Figure 13, B and C). 7 of 19 cell cycle-related genes and 5 of 13 proliferation-related genes were upregulated at least 2-fold (Supplemental Figure 13, D and E), implying that these cell cycle- and proliferation-related genes might play important roles in the regulation of heart regeneration in a Mettl3-m^6^A dependent manner. Among these genes, qPCR validation further confirmed that *Mis12*, *Fgf16*, and *Six5* were significantly upregulated during heart regeneration (Supplemental Figure 13F). To further explore the effects of these three genes on the proliferation of cardiomyocytes, H9c2 cells transfected with siRNAs were subjected to flow cytometry. We found that the silencing of *Fgf16*, rather than *Mis12* and *Six5*, significantly induced G1-phase cell cycle arrest (Supplemental Figure 13, G-J). Importantly, *Fgf16* silencing also suppressed the proliferation of primary cardiomyocytes isolated from neonatal mice, which was demonstrated by double staining with Ki67 and cTnT (Figure 8, A and B). Moreover, the decreased proliferation of primary cardiomyocytes induced by *Fgf16* silencing was further confirmed by nuclear EdU incorporation (Figure 8, C and D). Consistent with previous reports (Hotta *et al*. 2008; Yu *et al*. 2016), our results imply that Fgf16 is vital for promoting cardiomyocyte proliferation in neonatal mice.

**Figure 8.**
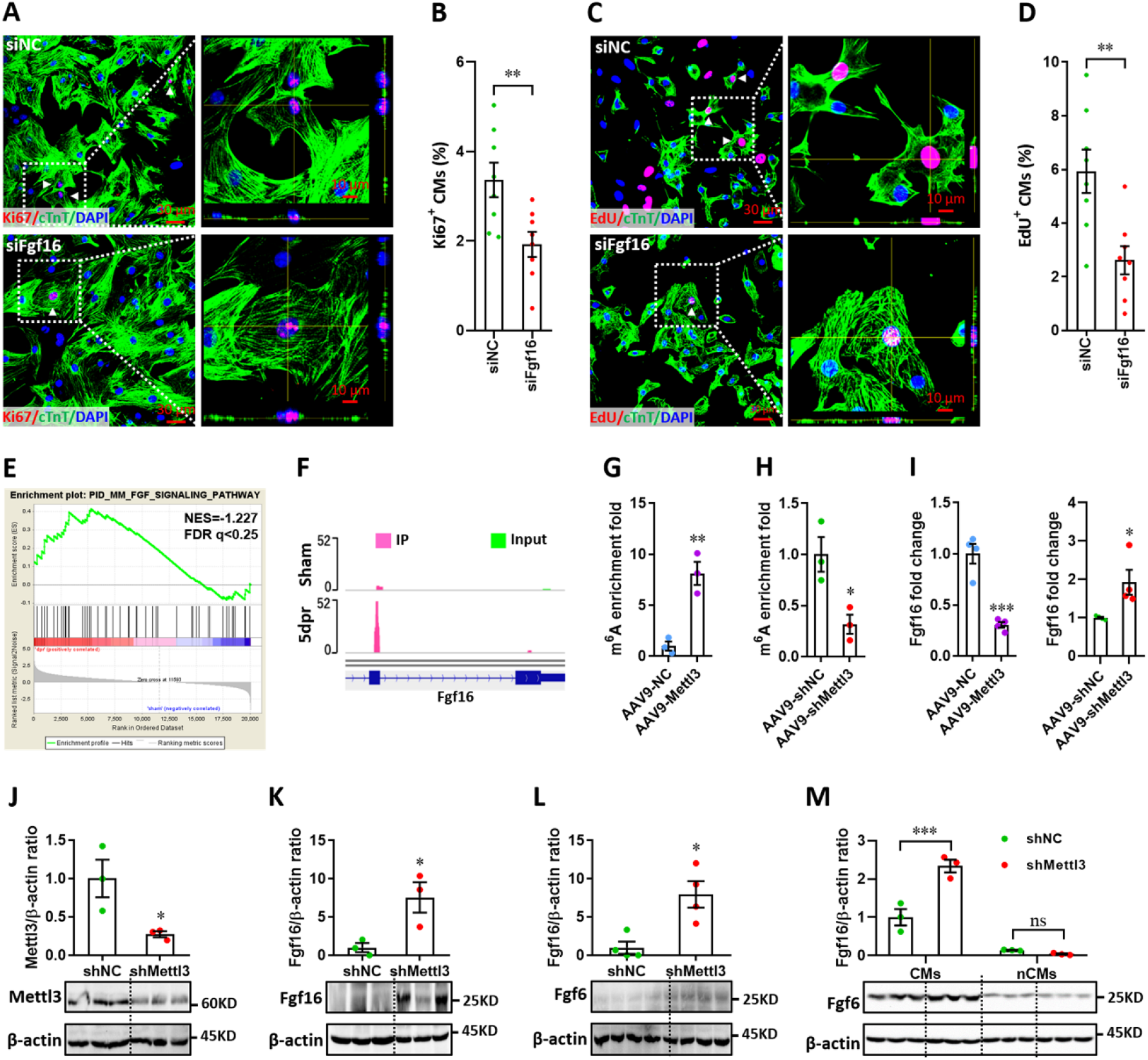
Mettl3-mediated m^6^A post-transcriptionally regulates Fgf16 during neonatal heart regeneration. (**A** and **B**) Primary cardiomyocytes were transfected with siFgf16 and siNC for 48 hr, followed by Ki67 (red) and cTnT (green) double staining. Representative images (A) and quantification (B) of Ki67^+^ cardiomyocytes are shown (*n*=8). Right panel (A), magnified Z-stack confocal images. (**C** and **D**) Primary cardiomyocytes were transfected with siFgf16 and siNC for 48 hr, followed by EdU incorporation assay. Representative images (C) and quantification (D) of EdU^+^ cardiomyocytes are shown (*n*=8). Right panel (C), magnified Z-stack confocal images. (**E**) GSEA analysis indicated the involvement of Fgf pathway in neonatal heart regeneration. (**F**) Representative IGV plots of mouse MeRIP reads for *Fgf16*. (**G** and **H**) MeRIP-qPCR quantification of m^6^A enrichment in *Fgf16* mRNA in Mettl3-overexpressing (G) and Mettl3-silencing (H) hearts at 5 dpr (*n*=3 hearts). (**I**) qPCR validation of *Fgf16* in the injured hearts with AAV9 virus injection at 5 dpr (*n*=4 hearts). (**J** and **K**) AAV9-shMettl3 viruses were injected into neonatal mice at p1, followed by apex resection at p3 and samples collection at 5dpr. Mettl3 knockdown (J, *n*=3 hearts) and Fgf16 expression (K, *n*=3 hearts) in neonatal hearts at 5 dpr were then validated by western blotting. (**L**) Validation of Fgf16 protein expression in homeostatic neonatal hearts at p8 after AAV9-shMettl3 injection at p1 (*n*=4 hearts). (**M**) Neonatal mice were injected with AAV9-shMettl3 virus at p1 and resected at p3. Fgf16 expression in primary cardiomyocytes (CMs) and non-cardiomyocytes (nCMs) isolated at 5 dpr were then validated by western blotting (*n*=3). All data are presented as the mean ± SEM, \**p*<0.05, \*\**p*<0.01, \*\*\**p*<0.001 versus control. *P* values were determined by 2-tailed Student’s *t* test (A-L), or by 2-way ANOVA with Dunnett’s multiple-comparison test (M).

GSEA revealed that the Fgf signaling pathway is related to neonatal heart regeneration (Figure 8E). Importantly, MeRIP-seq showed a great increase in the m^6^A peaks of *Fgf16* mRNA during neonatal heart injury (Figure 8F), implying the involvement of the m^6^A modification on *Fgf16* mRNA during heart regeneration. To further determine whether and how m^6^A modification regulates *Fgf16* mRNA levels, we performed m^6^A-RIP-qPCR and found that m^6^A enrichment in *Fgf16* was remarkably upregulated by *Mettl3* overexpression (Figure 8G) but suppressed by *Mettl3* silencing (Figure 8H) in the regenerating heart at 5 dpr. However, *Fgf16* mRNA expression at 5 dpr was greatly suppressed by *Mettl3* overexpression (Figure 8I). As expected, *Mettl3* silencing significantly increased the mRNA levels of *Fgf16* (Figure 8I). To further explore the effects of Mettl3 on Fgf16 expression in neonatal heart, western blotting was used to check the protein levels of Fgf16. Our data showed that Mettl3 knockdown (Figure 8J) significantly increased the production of Fgf16 protein in the regenerating heart at 5 dpr (Figure 8K). Mettl3 knockdown also increased the protein levels of Fgf16 at p8 (equal to 5 dpr in the injured heart) in the uninjured neonatal heart, although the expression levels were lower than that in the injured heart at 5 dpr (Figure 8L). Therefore, these findings suggest that Mettl3-mediated m^6^A negatively regulates the post-transcriptional levels of *Fgf16* during heart injury.

To further determine the cell types primarily expressed and secreted FGF16 in the heart, we examined the expression pattern of Fgf16 in cardiomyocytes (CMs) and non-cardiomyocytes (nCMs) during heart regeneration. After injection of AAV9 viruses at p1, CMs and nCMs were isolated from hearts in neonatal mouse at 5 dpr according to the method previously reported (Sander *et al*. 2013; Dorn *et al*. 2019). Isolated CMs and nCMs were directly subjected to protein extraction and western blotting assay. As shown in Figure 8M, Fgf16 expressions in CMs were greatly increased by Mettl3 deficiency compared with control hearts during regeneration. However, the expression of Fgf16 in nCMs was decreased in the Mettl3-deficient hearts compared with controls. Importantly, Fgf16 expression level in CMs are significantly higher than that in nCMs during heart regeneration regardless of Mettl3 deficiency or not (Figure 8M). Therefore, these findings indicate that the great changes in the m^6^A modification and expression of Fgf16 in the Mettl3-deficient hearts during regeneration dominantly results from CMs rather than nCMs.

To further address the effects of m^6^A modification on *Fgf16* expression, we constructed reporter genes by fusing luciferase to wild-type Fgf16 coding region with the m^6^A motif (GGACU) identified by MeRI-seq (Figure 8F) or to mutant Fgf16 (ΔFgf16) in which the m^6^A motif was replaced by GGGCU (Supplemental Figure 14A). The relative luciferase activity and mRNA level of the wild-type and mutant Fgf16-fused reporter genes were compared in H9c2 cells. For the wild-type reporter gene, both the luciferase activity and mRNA levels were greatly suppressed by *Mettl3* overexpression, and this suppression was significantly blocked by Mettl3 catalytic mutant (ΔMettl3) and the mutation of m^6^A consensus sequence in *Fgf16* (ΔFgf16), respectively (Supplemental Figure 14, B and C). Taken together, these findings suggest that the levels of *Fgf16* mRNA is negatively controlled by Mettl3-mediated m^6^A modification.

### Ythdf2 is involved in Mettl3-mediated downregulation of Fgf16

It has been demonstrated that the m^6^A methylation sites can be recognized by “readers”, and therefore impacting the fate of the target mRNA (Shi *et al*. 2019). Among the popular m^6^A “readers”, cytoplasmic Ythdf2 facilitates decay of its target transcripts with m^6^A modification under normal and stress conditions (Wang *et al*. 2014a; Du *et al*. 2016; Ivanova *et al*. 2017). By contrast, Ythdf1, the other m^6^A reader, promotes translation of target transcripts (Wang *et al*. 2015; Shi *et al*. 2018; Shi *et al*. 2019). To determine the potential involvement of these two representative m^6^A readers in Fgf16 regulation, we transfected primary cardiomyocytes with siRNAs and examined the expression of *Fgf16*. Silencing of *Ythdf2* rather than *Ythdf1* elevated the expression of *Fgf16* up to 6-fold in primary cardiomyocytes (Supplemental Figure 14, D and E). Moreover, we performed RNA decay assay and found that relative *Fgf16* mRNA level was increased in *Ythdf2*-deficient cardiomyocytes in comparison to control cells after Actinomycin-D treatment for 3 to 6 hr (Supplemental Figure 14F), indicating the decreased degradation of *Fgf16* mRNA. These data suggest that Ythdf2 may induce decay of *Fgf16* mRNA tagged by m^6^A, thereby negatively regulating *Fgf16* expression. In order to verify whether Ythdf2 participates in m^6^A modification of *Fgf16* mRNA, RIP-qPCR was used to investigate the interaction between *Fgf16* mRNA and Ythdf2. Our data showed that Ythdf2 remarkably enriched in *Fgf16* mRNA in normal primary cardiomyocytes, whereas this relative enrichment was significantly suppressed in *Mettl3*-deficient cardiomyocytes (Supplemental Figure 14G).

To further elucidate whether Ythdf2-regulated expression of *Fgf16* depends on m^6^A modification, wild-type and *Ythdf2*-deficient (Supplemental Figure 14H) H9c2 cells were co-transfected with wild-type or mutant Fgf16 reporter genes, respectively. Our data showed that the relative luciferase activity of wild-type reporter gene was greatly increased by *Ythdf2* knockdown. For mutant Fgf16 reporter plasmid, *Ythdf2* knockdown did not alter the relative luciferase activity. However, the mutant Fgf16 reporter plasmid showed increased luciferase activity in normal cells (siNC) compared with the wild-type reporter plasmid (Supplemental Figure 14I). These results suggest that Ythdf2 is involved in the Mettl3-mediated downregulation of Fgf16 by inducing decay of *Fgf16* mRNA.

### Effects of Fgf16 mutant on neonatal heart regeneration

Finally, we investigated the effects of Fgf16 mutant (ΔFgf16) on neonatal heart regeneration. To overexpress wild-type and mutant Fgf16 in cardiomyocytes, AAV9-Fgf16 and AAV9-ΔFgf16 viruses (Figure 9A) were injected into neonatal mice at p1, followed by apex resection at p3 and sample collection at the indicated time points (Figure 9B). AAV9-NC virus was used as negative control. The expression of target gene was determined by qPCR and western blotting at 5 dpr. Our data showed that the mRNA level of *Fgf16* in neonatal heart was slightly elevated by AAV9-Fgf16 virus. However, AAV9-ΔFgf16 remarkably increased the expression of *Fgf16* mRNA compared with AAV9-Fgf16 group (Figure 9C). To determine whether ΔFgf16 influences cardiomyocyte proliferation during heart regeneration, immunofluorescent staining was performed in cardiac tissues at 5 dpr using different proliferating markers including Ki67, pH3 and Aurora B. Ki67 staining showed that there was no significant difference in the proliferation of cardiomyocytes in cardiac apex in AAV9-Fgf16 group in comparison to AAV9-NC group. However, AAV9-ΔFgf16 significantly increased the percentage of Ki67^+^ cTnT^+^ cells compared with AAV9-Fgf16 group (Figure 9, D and E). The percentage of pH3^+^ cTnT^+^ cells was also significantly elevated by AAV9-ΔFgf16 compared with AAV9-Fgf16, although AAV9-Fgf16 only slightly promoted the percentage of pH3^+^ cTnT^+^ cells in comparison to AAV9-NC (Figure 9, F and G). Moreover, increased percentage of AurkB^+^ cardiomyocytes was also detected in cardiac apex in AAV9-ΔFgf16 group compared with AAV9-Fgf16 group (Figure 9, H and I). These findings strongly revealed that the mutation of m^6^A consensus sequence in *Fgf16* (ΔFgf16) can attenuate m^6^A/Ythdf2-mediated RNA decay during heart regeneration, thereby promoting cardiomyocyte proliferation. As expected, trichrome staining results further revealed a decreased fibrotic scar size at 14 dpr in AAV9-ΔFgf16 group compared with AAV9-Fgf16 group (Figure 9, J), indicating the accelerated heart regeneration. Consistent with these results, cardiac function analysis revealed that there was a higher left ventricular systolic function in AAV9-ΔFgf16 group compared with AAV9-Fgf16 group (Figure 9, K-M). Taken together, these results suggest that the mutation of m^6^A consensus sequence in *Fgf16* (ΔFgf16) can accelerate postnatal heart regeneration by promoting cardiomyocyte proliferation.

**Figure 9.**
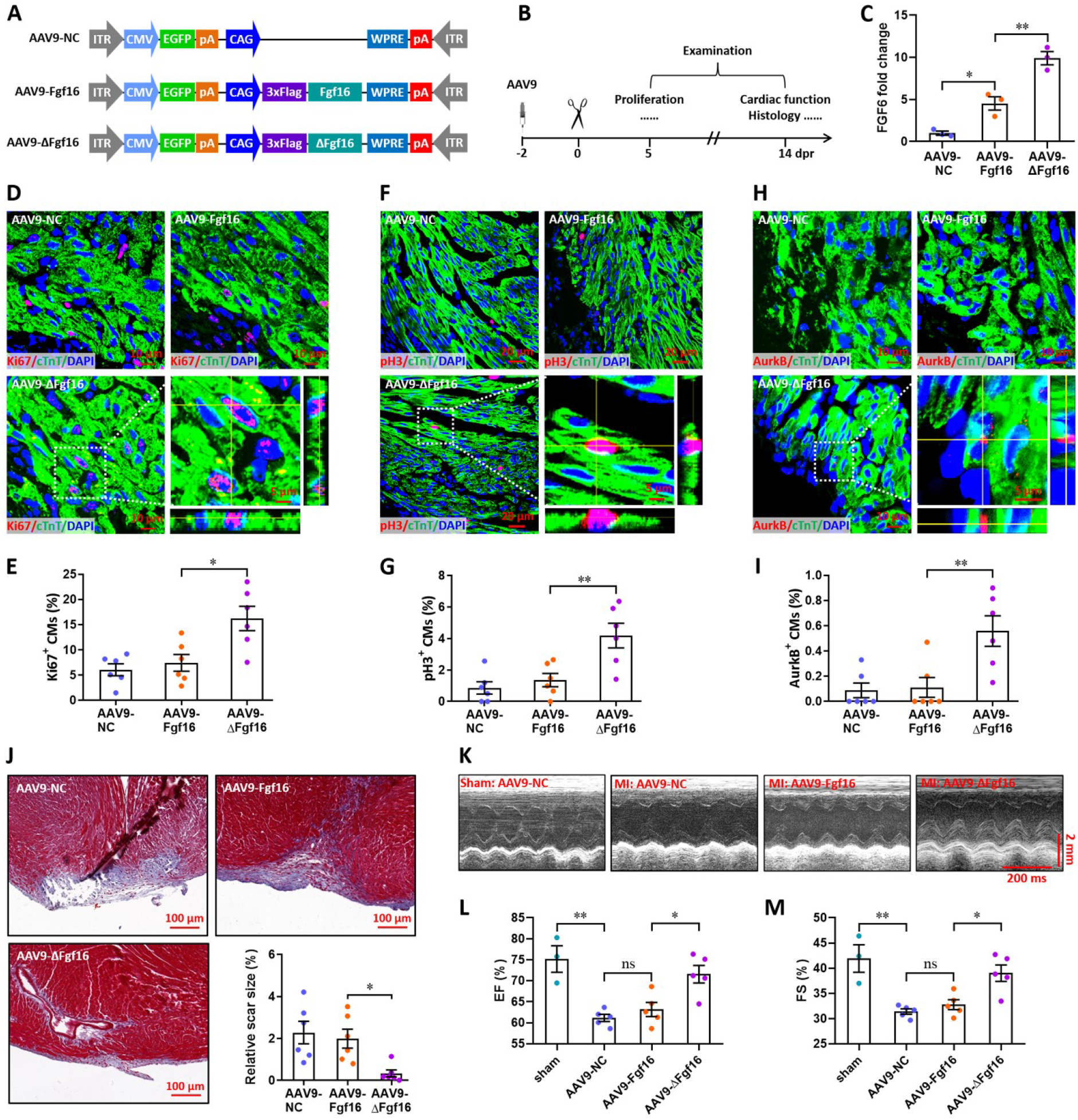
Mutation of the consensus sequence of m^6^A modification in *Fgf16* mRNA promotes heart regeneration. (**A**) Schematic of AAV9 vectors for the expression of negative control (AAV9-NC), wild-type Fgf16 (AAV9-Fgf16), and mutant Fgf16 (AAV9-ΔFgf16). (**B**) Schematic of AAV9 virus injection, apex resection, and sample collection in neonatal mice. (**C**) qPCR validation of *Fgf16* expression in neonatal hearts at 5 dpr (*n*=3 hearts). (**D** and **E**) Representative images (D) and quantification (E) of Ki67^+^ cardiomyocytes in apical ventricle at 5 dpr (*n*=6 hearts). Right lower panel (D), magnified Z-stack confocal image of Ki67^+^ cardiomyocyte. (**F** and **G**) Representative images (F) and quantification (G) of pH3^+^ cardiomyocytes in apical ventricle at 5 dpr (*n*=6 hearts). Right lower panel (F), magnified Z-stack confocal image of pH3^+^ cardiomyocyte. (**H** and **I**) Representative images (H) and quantification (I) of AurkB^+^ cardiomyocytes in apical ventricle at 5 dpr (*n*=6 hearts). Right lower panel (H), magnified Z-stack confocal image of AurkB^+^ cardiomyocyte. (**J**) Representative masson’s trichrome staining images of cardiac apex and quantification of scar size in hearts at 14 dpr (*n*=6 hearts). (**K**-**M**) Representative images of M-model echocardiography (K) and quantification of LVEF (L) and LVFS (M) are shown at 14 dpr (*n*=3 mice for sham, and 5 mice for each injured group). All data are presented as the mean ± SEM, \**p*<0.05, \*\**p*<0.01 versus control, ns means no significant difference. *P* values were determined by 1-way ANOVA with Dunnett’s multiple-comparison test.

## Discussion

In this study, we provide compelling in vitro and in vivo evidence demonstrating the important role of Mettl3 in cardiomyocyte proliferation and heart regeneration in mice upon injury. We identified Mettl3 as a key myocardial methylase that regulates cardiac m^6^A and provided a novel characterization of Mettl3-mediated m^6^A modification during heart regeneration in mice. Mettl3 deficiency increases cardiomyocyte proliferation in the early stage of injury and accelerates heart regeneration in postnatal mice (Figs. 2 to 4). However, overexpression of Mettl3 attenuated cardiomyocyte proliferation, heart regeneration and cardiac function (Figure 5). Our findings reveal that the Mettl3-mediated m^6^A modification of myocardial mRNAs plays an important role in regulating neonatal heart regeneration, demonstrating novel therapeutic potential for Mettl3-mediated m^6^A modification in heart regeneration.

Mechanistically, our work demonstrates a critical role for m^6^A modification in the regulation of mammalian heart regeneration upon injury. Indeed, several studies have recognized the importance of m^6^A modification in tissue and/or cell regeneration. The sciatic nerve injury elevates the levels of m^6^A-tagged transcripts encoding many regenerative genes and protein translation machinery components in adult mouse dorsal root ganglion, indicating a critical epitranscriptomic mechanism for promoting injury-induced axon regeneration in the adult mammalian nervous system (Weng *et al*. 2018). Moreover, expression of *Ythdf2* in hematopoietic stem cells (HSCs) facilitates the decay of the m^6^A-modified Wnt target gene mRNAs, thereby suppressing the Wnt signaling pathway. However, deletion of *Ythdf2* expands the number of HSCs and increased the regenerative capacity of HSCs under stress conditions, suggesting that Ythdf2-mediated m^6^A-dependent mRNA clearance facilitates hematopoietic stem cell regeneration (Wang *et al*. 2018b). A recent study revealed that Fto, an m^6^A demethylase, plays a critical role in cardiac contractile function during homeostasis, remodeling, and regeneration in an m^6^A-dependent manner (Mathiyalagan *et al*. 2019). These findings are consistent with our data and reveal that m^6^A modification is vital for the regulation of tissue regeneration.

m^6^A is added to mRNAs by a methyltransferase complex including Mettl3, Mettl14, and Wilm’s tumor 1-associating protein (Wtap), and among these proteins, Mettl3 was identified to function as a catalytic subunit (Bokar *et al*. 1997; Liu *et al*. 2014). In mammals, Mettl3 affects cell division, differentiation, reprogramming, and embryonic development. Knockout of *Mettl3* in mouse embryonic stem cells impairs differentiation and *Mettl3*^-/-^ mice are embryonic lethal (Geula *et al*. 2015). Consistent with these data, previous studies have demonstrated that Mettl3-mediated m^6^A modification is critical for cerebellar development (Wang *et al*. 2018a), HSC differentiation (Vu *et al*. 2017), and acute myeloid leukemia maintenance (Barbieri *et al*. 2017). These previous studies suggest an important role of Mettl3-mediated m^6^A mRNA methylation in physiology and pathology. In this study, *Mettl3* knockdown promoted cardiomyocyte proliferation and accelerated heart regeneration in postnatal mice in an m^6^A-dependent manner, whereas overexpression of *Mettl3* was sufficient to suppress cardiomyocyte proliferation, heart regeneration and cardiac function (Figures 2-5). Our data suggest that Mettl3-mediated m^6^A negatively regulates heart regeneration in neonatal mice. Consistent with our data in neonatal mice (Figures 4 and 5), previous study reported that *Mettl3* overexpression promotes cardiomyocyte hypertrophy in adult mice, but deficiency in *Mettl3* abolishes the ability of cardiomyocytes to undergo hypertrophy in response to stimulation (Dorn *et al*. 2019). Additionally, controversial functions of m^6^A modification were also detected in the differentiation and reprogramming of stem cells. For instance, *Mettl3* knockdown reduced self-renewal abilities in mouse embryonic stem cells (Wang *et al*. 2014b). However, a later study revealed a crucial role for *Mettl3* in embryonic priming from naïve pluripotency toward a more differentiated lineage (Geula *et al*. 2015). A reasonable explanation for the different functions of Mettl3 might be attributed to its multifaceted cellular localization, stability and translational modification under different conditions (Shi *et al*. 2019). In contrast with injured animal model, our data showed that Mettl3 gain and loss of function dose not significantly influence cardiomyocyte proliferation and cardiac function in the homeostatic postnatal mice without injury stresses (Supplemental Figures 4, 5, and 9). The different effects of Mettl3 between homeostatic and injured hearts might result from the robust changes in m^6^A modification during heart injury and regeneration. In agreement with this hypothesis, greatly increases in the levels of m^6^A modification and Mettl3 expression were indeed observed in the regenerating neonatal hearts at 5 dpr compared with sham hearts (Figure 1).

Using RNA-seq and MeRIP-seq, we identified Fgf16 as one of downstream targets of Mettl3-mediated m^6^A modification. This finding was further verified by MeRIP-qPCR and luciferase reporter gene assays. Mechanistically, our data demonstrate that Mettl3 epigenetically silences *Fgf16* in cardiomyocytes through an m^6^A-Ythdf2-dependent mechanism (Figures 8 and Supplemental Figure 14). Fgf16 belongs to the Fgf9 subfamily and is the only cardiac-specific Fgf family member. A previous study has reported the predominant expression of Fgf16 in cardiomyocytes and a significant decrease in the proliferation of embryonic cardiomyocytes in *Fgf16*^-/-^ mice (Hotta *et al*. 2008). Consistently, Fgf16 has been demonstrated to be required for embryonic heart development and cardiomyocyte replication (Lu *et al*. 2008). In addition, a recent study has shown that cardiac-specific overexpression of Fgf16 via AAV9 in *Gata4*-deficient hearts rescued cryoinjury-induced cardiac hypertrophy, promoted cardiomyocyte replication and improved heart function after injury (Yu *et al*. 2016). In line with these previous studies, our results show that *Fgf16* knockdown leads to a significant decrease in the proliferation levels of primary cardiomyocytes isolated from neonatal mice (Figure 8). In addition, the overexpression of mutant Fgf16 (mutation of m^6^A consensus sequence) can significantly accelerate postnatal heart regeneration by promoting cardiomyocyte proliferation compared with wild-type Fgf16 (Figure 9). Given that there is a higher level of m^6^A modification during heart regeneration (Figure 1), it is reasonable that wild-type *Fgf16* mRNA maintains a relative lower level compared with mutant *Fgf16*, due to the m^6^A/Ythdf2-induced mRNA decay.

It is well known that heart is a heterogeneous organ and consists of several different cardiac cell types including cardiomyocytes, fibroblasts, endothelial cells and others. Among them, cardiomyocyte is the largest cell type in whole heart and has the highest percentage in all heart cells. Particularly, ventricular regions contain 49.2% cardiomyocytes, 21.2% mural cells, 15.5% fibroblasts, 7.8% endothelial cells and 5.3% immune cells (Litvinukova *et al*. 2020). In the present study, ventricular regions rather than entire hearts were used for RNA-seq and MeRIP-seq analysis. Therefore, it is reasonable that the greatly changed genes identified by RNA-seq and changed m^6^A modification identified by MeRIP-seq during heart regeneration are likely to mainly result from cardiomyocytes. In line with this idea, the expression of Fgf16, the downstream target of Mettl3-mediated m^6^A modification identified by the conjoint analysis of MeRIP-seq and RNA-seq, was dominantly expressed in CMs rather than nCMs regardless of injury stimulation (Figure 8M).

Although the m^6^A modification is a part of the larger field of RNA epigenetics, yet its function during heart regeneration remains elusive. In this study, we demonstrate that Mettl3 post-transcriptionally reduces *Fgf16* mRNA levels through an m^6^A-Ythdf2-dependen pathway, thereby restricting cardiomyocyte proliferation during heart regeneration in mice. Our data exemplify the pivotal role of m^6^A epitranscriptomic changes in heart regeneration. Targeting the Mettl3-mediated m^6^A modification of Fgf16 may represent a promising therapeutic strategy for promoting the proliferation of cardiomyocytes in mammals. Our data on m^6^A-modification in hearts further our understanding of the mechanism of heart regeneration and provide innovative therapeutic interventions.

## Materials and Methods

### Cell culture and treatment

H9c2 cells (rat cardiomyocyte cell line) were purchased from FuDan IBS Cell Center (FDCC, Shanghai, China) and were cultured in high glucose DMEM (Corning, USA) with 10% fetal bovine serum (Gibco, USA), l µg/mL streptomycin (Gibco, USA), and 1 U/mL penicillin streptomycin (Gibco, USA), at 37℃ in a 5 % CO_2_ incubator. Culture medium was replaced every 2 to 3 days and cells were passaged when they reached 80% confluence. Primary cardiomyocytes were isolated from the hearts of neonatal mice as previously described (Nakada *et al*. 2017). To knock down or overexpress target genes in cardiomyocytes, several methods including siRNAs, lentivirus and adeno-associated virus 9 were used as described below.

### Knockdown and overexpression of target genes in H9c2 cells

For Mettl3 knockdown in H9c2 cells, the specific siRNA for Mettl3 (siMettl3) and negative control (siNC) were synthesized by RIBOBIO (Guangzhou, China). Cells were transfected with siMettl3 to knock down the expression of Mettl3. The knockdown of other target genes was also induced by siRNA transfection. Gene silencing was achieved by transfection of predesigned siRNA duplexes (Supplementary file 1) designed and synthesized by RIBOBIO (Guangzhou, China).

The coding sequence of Mettl3 was amplified from pcDNA3/Flag-METTL3 (Addgene, #53739) and inserted into the lentiviral plasmid pLOXCMV (Qi *et al*. 2015) to generate pLOX-Mettl3 overexpression plasmid. Viral particles were packaged in HEK 293T cells and used to infect H9c2 cells as previously described (Qi *et al*. 2015). The infected H9c2 cells were selected by puromycin and expanded to form a stable sub-line.

### Adeno-associated virus 9 (AAV9) production

The full-length Mettl3 coding sequences were amplified from the pcDNA3-Mettl3 plasmid (Addgene, #53739), and cloned into AAV serotype-9 expressing plasmid to overexpress Mettl3 (AAV9-Mettl3). AAV9-NC (virus packaged with empty plasmid) served as control for AAV9-Mettl3. In addition, Mettl3 specific shRNA or negative control (RiboBio, Guangzhou, China) were cloned into AAV9-shRNA expressing plasmid to generate AAV9-shMettl3 or AAV9-shNC plasmids, respectively. To overexpress wild-type Fgf16 and mutant Fgf16 (ΔFgf16) in which the adenosine bases (GGACT) in m^6^A consensus sequences were replaced by guanine (GGGCT), both wild-type and mutant Fgf16 coding sequences were constructed into AAV9 plasmids (Supplemental Figure 15A) for AAV9 virus generation. AAV9 viruses were packaged and produced using the AAV Helper-Free System (DongBio.Co.Ltd, Shenzhen, China). For AAV9 virus delivery *in vivo*, viruses were subcutaneously injected into neonatal mice at a dose of 1×10^11^ V.G./mouse at postnatal day 1 (p1). The schematic of AAV9 virus injection can be found in Figures 3-7, and 9, as well as in Supplemental Figures 2, 4-6 and 9. Primary cardiomyocytes were infected with AAV9 virus at a dose of 5×10^10^ V.G./well (24 well plate).

### Mettl3 knockdown and overexpression in primary cardiomyocytes

The isolation of primary cardiomyocytes was performed as previous described (Nakada *et al*. 2017). Briefly, fresh neonatal mice (within 2 days after birth) hearts were harvested and immediately fixed in 4% PFA/PBS at 4 ℃ for 4 hours. Hearts were subsequently incubated with collagenase IV (2.4 mg/ml, Sigma) and II (1.8 mg/ml, Sigma) for 12 hours at 37°C. The supernatant was collected and spun down in 500 rpm for 2 minutes to yield the cardiomyocytes. The hearts were then minced to smaller pieces and the procedure was repeated until no more cardiomyocytes were dissociated from the hearts. Knockdown and overexpression of target genes in primary cardiomyocytes was induced by siRNAs transfection and AAV9-mediated expression, respectively. Sequences of siRNAs used in this study are listed in Supplementary file 1.

### In vitro cell proliferation assay

For cell proliferation assay, both primary cardiomyocytes and H9c2 cell line were incubated in 24 well plates with different treatments. DNA synthesis were then analyzed by 5-Ethynyl-2’-deoxyuridine (EdU) labelling, using Cell-Light™ EdU Apollo®567 In Vitro Imaging Kit (RiboBio, Guangzhou, China) according to the manufacturer’s instructions. At least 7 images were randomly taken per well using a Zeiss LSM 700 laser confocal microscope (Carl Zeiss). The population of EdU^+^ cells was determined by counting at least 500 cells per well. The EdU^+^ cells were quantified as the percentage of total cells. Moreover, proliferation of primary cardiomyocytes was also performed by Ki67 (Abcam, ab16667, 1:250) staining. In addition, cell numbers were analyzed using the Enhanced Cell Counting Kit-8 (CCK-8, Beyotime Biotechnology, China) according to the manufacturer’s instructions, as previously described (Zhou *et al*. 2018).

### Animals and heart injury models

All animal studies were approved by the Institutional Animal Care and Use Committee of Jinan University and conducted in accordance with the ARRIVE guidelines (Percie du Sert *et al*. 2020). In vivo experiments and animal management procedures were performed according to the NIH Guide for the Care and Use of Laboratory Animals. Both male and female C57BL/6J mice were used for neonatal heart injury experiments. Apical resection surgeries were performed on neonatal mice at postnatal day 3 (p3) and p7, respectively, as described previously (Porrello *et al*. 2011). Hearts with or without EdU injection were collected at the indicated time points for further analysis. For *in vivo* knockdown or overexpression of target genes, AAV9 viruses were injected at p1, followed by apex resection (at p3 and p7) and sample collection at the indicated time points. In addition, myocardium infarction (MI) injury in adult male mice (8 weeks) were induced by permanent ligation of the left anterior descending artery (LAD) as previously described (Wu *et al*. 2021). In cases when anaesthesia was required, mice were anaesthetized by oxygen and 2% isoflurane as previously reported (Scott *et al*. 2021). Animals were euthanized by dissecting the diaphragm under isoflurane anaesthesia, after which organs were harvested.

### Measurement of m^6^A Level

Total RNA was isolated using RNeasy Kit (Qiagen, Valencia, CA, USA) from heart tissue according to the protocol of the manufacturer. The mRNA was then further purified using the Dynabeads™ mRNA Purification Kit (ThermoFisher). The m^6^A levels in mRNA were detected by colorimetric ELISA assay with the EpiQuikTM m^6^A RNA Methylation Quantification kit (Epigentek, P-9005). Measurements were carried out in triplicate for each sample and 5 hearts were used for each group.

### Echocardiography

After AAV9 virus injection, animals with or without heart injury were subjected to echocardiography analysis at indicated time points. Cardiac function was evaluated using the Vevo® 2100 ultrasound system (Visualsonics, Toronto, Canada) equipped with a high-frequency (30 MHz) linear array transducer, as described previously (Wu *et al*. 2021).

### Histology

Mice were sacrificed and weighed to obtain total body weight (BW) at indicated time points. The heart was then harvested and weighed to obtain heart weight (HW) and HW/BW ratio. Harvested hearts were fixed in 4% paraformaldehyde (PFA)/PBS solution overnight at room temperature, dehydrated in an ethanol series, and then processed for paraffin embedding. Paraffin sections were cut in 5 μm thickness. Sections were subjected to Masson’s trichrome staining according to standard procedures. Fibrotic scar size was measured using the CaseViewer version 2.1 software (3DHISTECH, Hungary). The whole images of hearts were captured by Leica M205FA stereo fluorescence microscope to visualize the apex regeneration at indicated time points.

### Immunofluorescence staining

Hearts were embedded in paraffin and cut in 5 μm sections, deparaffinized, rehydrated and antigen retrieval. Sections were permeabilized with 0.5% Triton X-100/PBS and then blocked with 5% goat serum (Jackson ImmunoResearch Laboratories, USA) for 1 hour at room temperature, and incubated with primary antibodies overnight at 4℃. After washing with PBS, sections were incubated with corresponding secondary antibodies conjugated to fluorescence for 1 hour at room temperature, followed by counterstaining with DAPI (Sigma). Primary antibodies are as follows: anti-Mettl3 (Abcam, ab195352, 1:500), anti-m^6^A (Synaptic Systems, 202111, 1:100), anti-Ki67 (Abcam, ab16667, 1:250), anti-phospho-Histone H3 (pH3) (CST, #53348S, 1:400), anti-aurora kinase B (AurkB) (Abcam, ab2254, 1:200,) anti-GFP (Proteintech, 50430-2-AP, 1:200), and anti-Cardiac Troponin T (cTnT) (ThermoFisher, MA512960, 1:200). Secondary antibodies used are following: Alexa Fluor 488 goat anti-mouse or anti-rabbit IgG (Jackson ImmunoLabs, 115-545-071 or 111-545-003, 1:200), and Cy3-conjugated Affinipure Goat anti-mouse or anti-rabbit IgG (Proteintech, SA00009-1 or SA00009-2, 1:300). The slides were imaged with fluorescence microscope (Leica Microsystems) or Zeiss LSM 700 laser confocal microscope (Carl Zeiss).

### Wheat germ agglutinin (WGA) staining

Following deparaffinized, rehydrated, slides were then incubated with WGA conjugated to Alexa Fluor 488 (Invitrogen, W11261, 5 μg/ml) for 10 minutes at room temperature. To quantify the cell size, 5 independent hearts per group (at least 300 cells) were captured near apex with laser-scanning confocal microscope (LSM 700, Zeiss). ZEN 2012 lite software (Zeiss) was used to quantify the size of each cell.

### EdU labeling assay in vivo

For EdU labeling experiments, neonatal mice were subcutaneously injected with 50 μl of a 2 mg/ml solution of EdU (RiboBio, Guangzhou, China) dissolved in sterile water. Hearts were embedded in Tissue-Tek optimal cutting temperature compound (OCT) (Sakura, USA) for frozen section (4 μm). Sections were rinsed three times in PBS and fixed in 4% parapormaldehyde for 30 minutes. After rinsing three times again, citrate antigen retrieval was performed as described above. Sections were then incubated with 2 mg/mL glycine solution for 10 minutes, permeabilized with 0.5% Triton X-100 in PBS for 10 minutes, and then rinsed with PBS once for 5 minutes. This was followed by incubation with Apollo®576 staining solution (1×) at room temperature for 30 minutes. Permeabilization was performed again with 0.5% Triton X-100 in PBS twice for 10 minutes. Sections were then rinsed with methanol for 5 minutes, washed with PBS once for 5 minutes, blocked with 5% goat serum for 1h, and followed by incubation with primary antibody against cTnT (ThermoFisher, MA512960, 1:200) overnight. The following day, incubation with anti-mouse secondary antibody conjugated to Alexa Fluor 488 (1:200 dilution, Jackson ImmunoResearch Laboratories, USA) was performed. Sections were washed three times in PBS, stained with DAPI for 10 minutes to label nuclei, and mounted in Antifade Mounting Medium. Images were captured by laser-scanning confocal microscope (LSM 700, Zeiss) and analyzed by ZEN 2012 software (Zeiss).

To analyze CMs proliferation at the indicated time points, EdU was injected 8 hours prior to heart collection. For EdU pulse-chase experiments, EdU was injected once every two days to label all proliferating CMs during the whole period of cardiac regeneration. The last injection was performed 8 hours prior to heart collection. Sham-operated mice underwent the same procedure without the apical resection.

### RNA-seq

For total RNA isolation, neonatal mice were subjected to cardiac apical resection at postnatal day 3 (p3). Hearts were then extracted in sham and injured neonatal mice at 5 days post-resection (dpr), respectively. Three ventricles per group were used for RNA-sequencing analysis. RNA preparation, library construction and sequencing on GBISEQ-500 platform was performed as previously described (Xin *et al*. 2017). After filtering the reads with low quality, clean reads were then obtained and mapped to the reference genome of mouse (GRCm38.p6) with HISAT (Kim *et al*. 2015). Genes expression level was quantified by a software package called RSEM (Li & Dewey 2011) and expressed as fragments per kilobase of exon per million fragments mapped (FPKM). Differential expressed (DE) genes were detected using NOISeq method (Tarazona *et al*. 2011) with Probability ≥ 0.8 and fold change (FC) of FPKM ≥ 2. Only those genes were considered for the differential expression analysis, which displayed FPKM ≥ 1 in either of the two samples under comparison. GO analysis was performed using online tool DAVID 6.8 (https://david.ncifcrf.gov/summary.jsp), and terms with *p-*value ≤ 0.05 were included. Differentially expressed gene heat maps were clustered by hierarchical clustering using cluster software (Eisen *et al*. 1998). Gene set enrichment analysis (GSEA) was performed to identify gene sets from signaling pathways that showed statistical differences between two groups by using GSEA software (http://software.broadinstitute.org/gsea/index.jsp).

### MeRIP-seq

Messenger RNAs isolated from neonatal hearts at 5dpr and sham-operated hearts were subjected to MeRIP-seq. Eight neonatal hearts were pooled with at least two biological duplicates for each group. MeRIP-Seq was performed by Cloudseq Biotech Inc. (Shanghai, China). Briefly, fragmented RNA was incubated with anti-m^6^A polyclonal antibody (Synaptic Systems, 202003) in IPP buffer for 2 hours at 4°C. The mixture was then immunoprecipitated by incubation with protein-A beads (Thermo Fisher) at 4°C for an additional 2 hours. Then, bound RNA was eluted from the beads with N6-methyladenosine (BERRY & ASSOCIATES, PR3732) in IPP buffer and then extracted with Trizol reagent (Thermo Fisher) by following the manufacturer’s instruction. Purified RNA was used for RNA-seq library generation with NEBNext® Ultra™ RNA Library Prep Kit (NEB). Both the input sample without immunoprecipitation and the m^6^A IP samples were subjected to 150 bp paired-end sequencing on Illumina HiSeq sequencer. For data analysis, Paired-end reads were harvested from Illumina HiSeq 4000 sequencer, and were quality controlled by Q30. After 3’ adaptor-trimming and the lower quality reads removing by cutadapt software (v1.9.3). First, clean reads of all libraries were aligned to the reference genome (UCSC MM10) by Hisat2 software (v2.0.4). Methylated sites on RNAs (peaks) were identified by MACS software. Differentially methylated sites were identified by diffReps. These peaks identified by both software overlapping with exons of mRNA were figured out and chosen by home-made scripts. Differential peak analyses of MeRIP-seq data sets were performed by using a modification of the exomeReak R/Bioconductor package to compare the ratio of the absolute number of MeRIP reads with nonimmunoprecipitation reads at a given peak between 2 conditions (Meng *et al*. 2014; Mathiyalagan *et al*. 2019). Both RNA-seq and MeRIP-seq can be accessed in the Sequence Read Archive (SRA) under accession numbers SRP224051.

### RNA extraction and quantitative Real-Time PCR (qPCR)

Total RNA was isolated using RNeasy Kit (Qiagen, Valencia, CA, USA) from cells or heart tissue according to the protocol of the manufacturer, respectively. Reverse transcription to cDNA was performed with 30 ng of total RNA, random primers, and SuperScript III Reverse Transcriptase (Roche, USA). The qPCR was performed using a Light Cycler 480 SYBR Green I Master (Roche, USA) and the MiniOpticon qPCR System (Bio-Rad, CA, USA). After denaturation for 10 min at 95 °C, the reactions were subjected to 45 cycles of 95 °C for 30 s, 60 °C for 30 s, and 72 °C for 30 s. GAPDH was used as the internal standard control to normalize gene expression using the 2^-ΔΔCt^ method. The sequences of the qPCR primers were listed in Supplementary files 2 and 3.

### Protein extracts and Western blotting

Tissues or cells for SDS-PAGE were lysed in RIPA buffer (Beyotime Biotechnology) containing protease inhibitors (Sigma). Protein concentration ware determined using the Bio-Rad Protein Assay (Bio-Rad Laboratories). 30 μg of protein were separated by SDA-PAGE, proteins were transferred onto PVDF membranes (Millipore), then blocked in 5% nonfat milk/TBS-Tween 20 and incubated with primary antibodies (dilution in TBST) overnight at 4℃. Membranes were then washed and incubated with corresponding second antibodies for 1 hour at room temperature. Bands were detected by chemiluminescence reagents (ThermoFisher Scientific). Primary antibodies used are following: anti-Mettl3 (Abcam, ab195352, 1:1000), anti-Mettl14 (Abcam, ab98166, 1:1000), anti-Fgf16 (Santa Cruz, sc-390547, 1:500), anti-GFP (CST, #2956, 1:1000), anti-Flag (CST, #14793, 1:1000), and anti-β-actin (Proteintech, 60008-1-Ig, 1:2000). Secondary antibodies used are following: goat-anti-mouse horseradish peroxidase (HRP)-conjugated antibody (CST, #7076, 1:3000) and goat-anti-rabbit horseradish peroxidase (HRP)-conjugated antibody (CST, #7074, 1:2000). Quantitation of the chemiluminescent signal was analyzed using Image-Pro Plus version 6.0 (Media Cybernetics, Bethesda, MD). The relative expression levels of target protein/β-actin were set as one. All blots derive from the same experiment and were processed in parallel. The uncropped blots are listed in the Source data.

### Luciferase reporter assay

The pCMV-Gaussia-Dura Luc and pTK-Red Firefly Luc plasmids from ThermoFisher were used to construct the dual-luciferase reporter plasmid (pGL-RF), which contained both a Gaussia luciferase (GL) and a red firefly luciferase (RF). In brief, a new combined DNA fragment (Gaussia Dura Luc-SV40pA-pTK-Red Firefly Luc-bGHpA, GL-SV40pA-pTK-RF-bGHpA) was synthesized by Shanghai Generay Biotech Co., Ltd (Shanghai, China). In the synthesized fragment, *XhoI* and *ClaI* restriction sites were inserted upstream and downstream of the termination codon of GL gene, respectively, to subclone Fgf16 coding sequences as below. The pTK-RF-bGHpA cassette from pTK-Red Firefly Luc plasmid was synthesized with replacement of *BamHI* restriction site between pTK promoter and RF gene with *XbaI*. The pCMV-Gaussia-Dura Luc plasmid was digested by *BamHI* and *NaeI* enzymes to remove the GL-bGHpA-SV40 promoter-Puromycin resistant gene (GL-bGHpA-SV40p-PuroR) fragment. The residual plasmid frame was then inserted by the synthesized combined fragment (GL-SV40pA-pTK-RF-bGHpA) using the EasyGeno Single Assembly Cloning kit (Tiangen Biotech, Beijing, China) to construct the dual-luciferase reporter plasmid (pGL-RF). The DNA sequence of pGL-RF plasmid can be found in Supplementary file 4. To constructed the wild-type Fgf16 reporter plasmid (pGL-Fgf16), the full-length of Fgf16 coding sequence (NM_030614.2) was amplified and subcloned into the pGL-RF plasmid linearized by digestion of *XhoI* and *ClaI* enzymes, thereby expressing the fusion protein of Gaussia-Fgf16. To make the mutant Fgf16 reporter plasmid (pGL-ΔFgf16), the adenosine bases in the m^6^A consensus sequence (GGACT) were replaced with guanine (GGGCT) using a KOD-Plus-Mutagenesis Kit (Toyobo). H9c2 cells seeded in 96 well plates were co-transfected with reporter plasmid and pcDNA3-Mettl3 (Addgene, #53739) (100 ng for each) using LipoFiter Liposomal Transfection Reagent (Hanbio Biotechnology). An Mettl3 catalytic mutant (AA395-398, DPPW to APPA) was generated using a KOD-Plus-Mutagenesis Kit (Toyobo) to generate pcDNA-ΔMettl3 as described previously (Li *et al*. 2019). Two days after transfection, luciferase activity was examined using the Pierce^TM^ Gaussia-Firefly Luciferase Dual Assay Kit (ThermoFisher Scientific) according to the manufacturer’s protocol. Relative luciferase activity was measured using a BioTek Synergy^TM^ 4 multimode microplate reader (BioTek Instruments). The activity of the Gaussia luciferase was normalized with that of Firefly luciferase. In addition, the mRNA expression levels of Gaussia luciferase were examined by qPCR as described above. Primers for Gaussia luciferase are as follows: Fw, 5’-ACC ACG GAT CTC GAT GCT GAC-3’; Re, 5’-ACT CTT TGT CGC CTT CGT AGG TG-3’. GAPDH was used as the internal standard control to normalize gene expression using the 2^-ΔΔCt^ method.

### MeRIP-qPCR and Ythdf2-RIP-qPCR

The m^6^A-RIP-qPCR and Ythdf2-RIP-qPCR were performed according to a protocol as described previously (Wang *et al*. 2018b) with some modification. In brief, total RNA was isolated from neonatal hearts injected with AAV9-Mettl3 virus. The mRNA was then further purified using the Dynabeads™ mRNA Purification Kit (ThermoFisher) and fragmented by the RNA Fragmentation Reagents (ThermoFisher). The fragmented mRNA was incubated with anti-m^6^A antibody (Synaptic Systems), anti-Ythdf2 antibody (Proteintech), or IgG in IPP buffer (150 mM NaCl, 10 mM TRIS-HCL and 0.1% NP-40) for 4 hr, followed by incubation with the eluted and blocked Dynabeads Protein A (ThermoFisher) for 2 hr at 4°C. The mRNA binding to beads was eluted and purified with RNeasy MinElute Spin columns (Qiagen), followed by cDNA synthesis. qPCR was then performed as above described. GAPDH was used the control. The m^6^A or Ythdf2 enrichment of Fgf16 was evaluated by 2^-ΔΔCt^ method. Primers used for RIP-qPCR in this study are as follows: RIP-Fw, 5’-GAC CAC AGC CGC TTC GGA AT-3’; RIP-Re, 5’-CGA TCC ATA GAG CTC TCC TCG C-3’.

### Measurement of Fgf16 mRNA stability

Primary cardiomyocytes were isolated from neonatal mice and treated with vehicle or Actinomycin-D (Sigma-Aldrich) at a concentration of 5 μM for 0, 3, and 6 hr, respectively. RNA was extracted at the indicated timepoints. qPCR analysis was then performed as described in “RNA extraction and quantitative Real-Time PCR (qPCR)” section. Actinomycin-D was used to inhibit global mRNA transcription, and the ratio of *Fgf16* mRNA in Actinomycin-D-treated cells relative to vehicle-treated cells (A/V ratio) was calculated to evaluate the stability of *Fgf16* mRNA.

### Statistics

All statistics were calculated using GraphPad Prism 8 Software. Among three or more groups, statistical analysis was performed using one-way or two-way ANOVA with Dunnett’s multiple comparisons post hoc tests. Comparisons between two groups were analyzed using unpaired and 2-tailed Student’s *t*-test. All data are presented as the mean ± SEM. A *p* value of less than 0.05 was considered statistically significant.

## Author Contributions

X.F.Q. designed the study. F.Q.J., J.X.C., W.Y.C., and W.L.Z. performed most experiments, and analyzed the data. C.Q.L. and Y.M.Z. contributed plasmids construction, animal and cellular experiments. G.H.S., H.L.H., R.J.H., H.Z., and K.S.P. provided valuable comments and suggestions. D.Q.C. and Z.Y.J. revised and edited the manuscript. X.F.Q. conceived of and supervised the study, and wrote the manuscript with helps from co-authors.

## Acknowledgments

This work was supported by grants from the National Key R&D Program of China (2017YFA0103302), the National Natural Science Foundation of China (82070257, 81770240, 81570222, 81670422, 81873517), the Guangdong Natural Science Funds for Distinguished Young Scholar (2014A030306011), the Guangdong Science and Technology Planning Project (2014A050503043), the New Star of Pearl River on Science and Technology of Guangzhou (2014J2200002), the Top Young Talents of Guangdong Province Special Support Program (87315007), the Young Taishan Scholars Program of Shandong Province (tsqn20161045), and the Research Grant of Key Laboratory of Regenerative Medicine, Ministry of Education, Jinan University (ZSYX-M-2019-00009, ZSYXM202004, and ZSYXM202104), China.

## Figures and figure legends

**Supplemental Figure 1.**
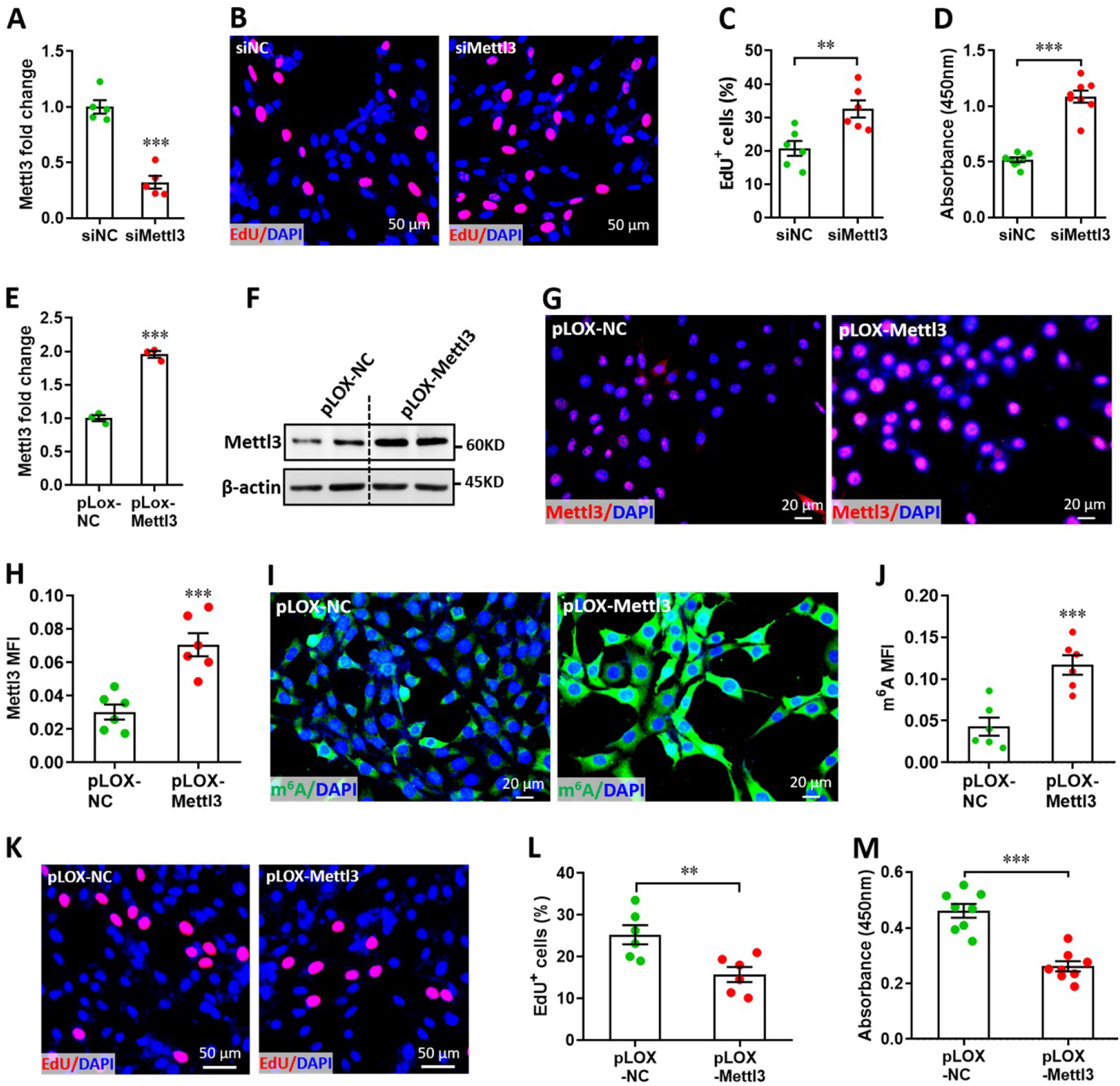
Effects of Mettl3 on the proliferation of H9c2 cells. (**A**) siMettl3-induced silence of *Mettl3* mRNA in H9c2 cells was confirmed by qPCR (*n*=3). (**B**-**D**) H9c2 cells were transfected with siMettl3 or siNC for 48 hr, followed by EdU labeling or cell counting examination. Representative images of EdU labeling (B) and quantification of EdU^+^ cells (C) are shown (*n*=6). Cell counting (D) was examined using CCK8 Cell Counting Kit (*n*=8). (**E** and **F**) Stable Mettl3 overexpression H9c2 cell line was established using lentivirus system. Mettl3 overexpression in mRNA and protein levels were evidenced by qPCR (E) and western blotting (F) respectively (*n*=3). (**G** and **H**) Representative images of immunofluorescence staining (G) and quantification of FoxO3 expression (H) are shown (*n*=6). (**I** and **J**) Immunofluorescence staining confirmed the increased m^6^A levels in the stable Mettl3 overexpression H9c2 cells. Representative images (I) and quantification (J) of m^6^A staining are shown (*n*=6). (**K**-**M**) Stable H9c2 cell lines screened by pLOX-Mettl3 and pLOX-NC were subjected to EdU labeling or cell counting examination. Representative images of EdU labeling (K) and quantification of EdU^+^ cells (L) are shown (*n*=6). Cell counting (M) was examined using CCK8 Cell Counting Kit (*n*=8). All data are presented as the mean ± SEM of three separate experiments, \*\**p*<0.01, \*\*\**p*<0.001 versus control. *P* values were determined by 2-tailed Student’s *t* test.

**Supplemental Figure 2.**
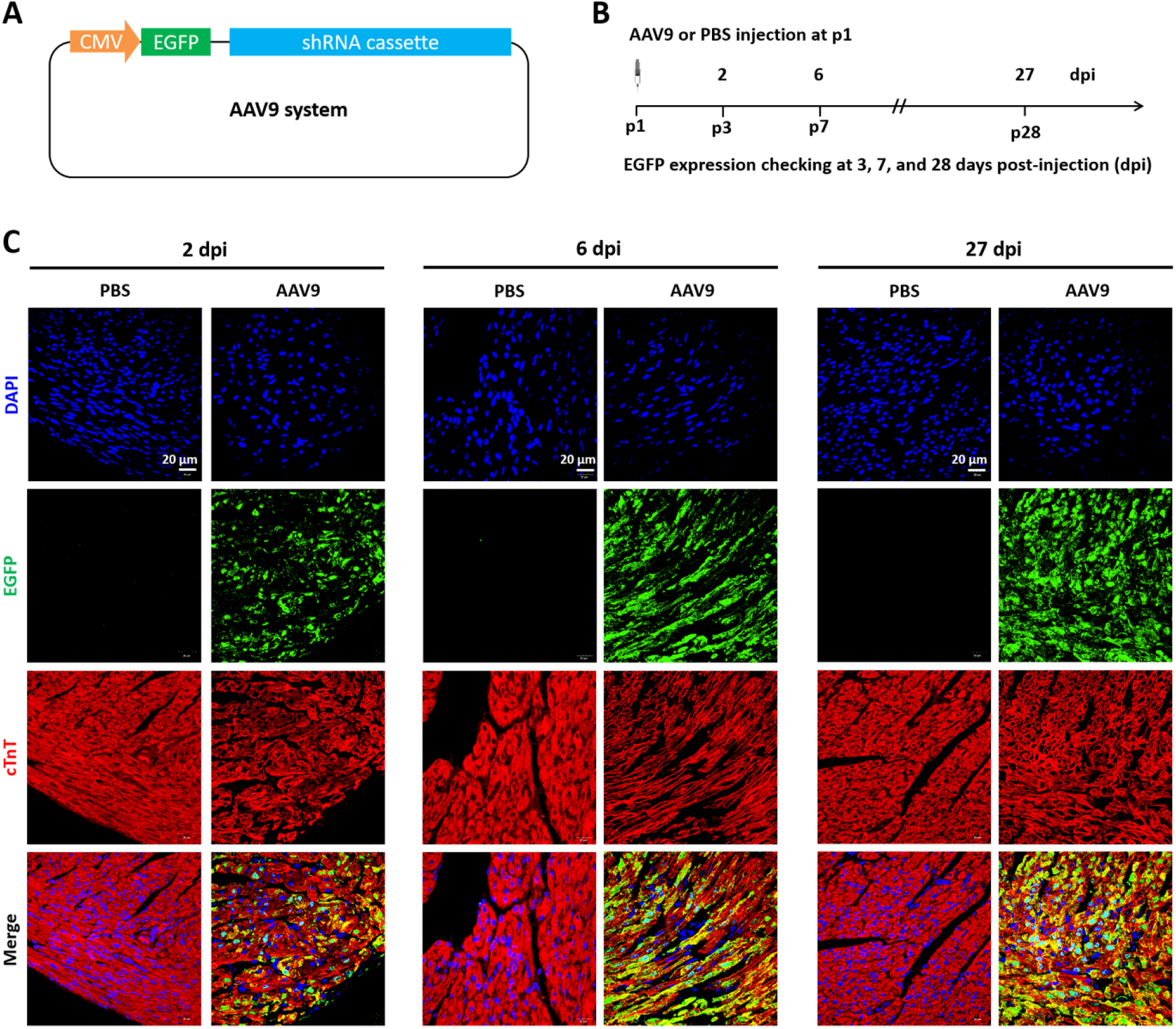
Time course of AAV9 system infection in neonatal mice heart. (**A**) A diagram of the AAV9 system harboring a reporter gene (EGFP) expression cassette. (**B**) Schematic of AAV9 virus injection experiment designed to verify the infective efficiency in neonatal hearts. AAV9 virus was injected at p1, followed by detection of reporter gene at 2-27 days post-injection (dpi) using immunofluorescent staining. (**C**) Representative images of EGFP expression (green) in heart at 2, 6, and 27 dpi. Red color (cTnT) and blue color (DAPI) denote cardiomyocytes and nucleus, respectively.

**Supplemental Figure 3.**
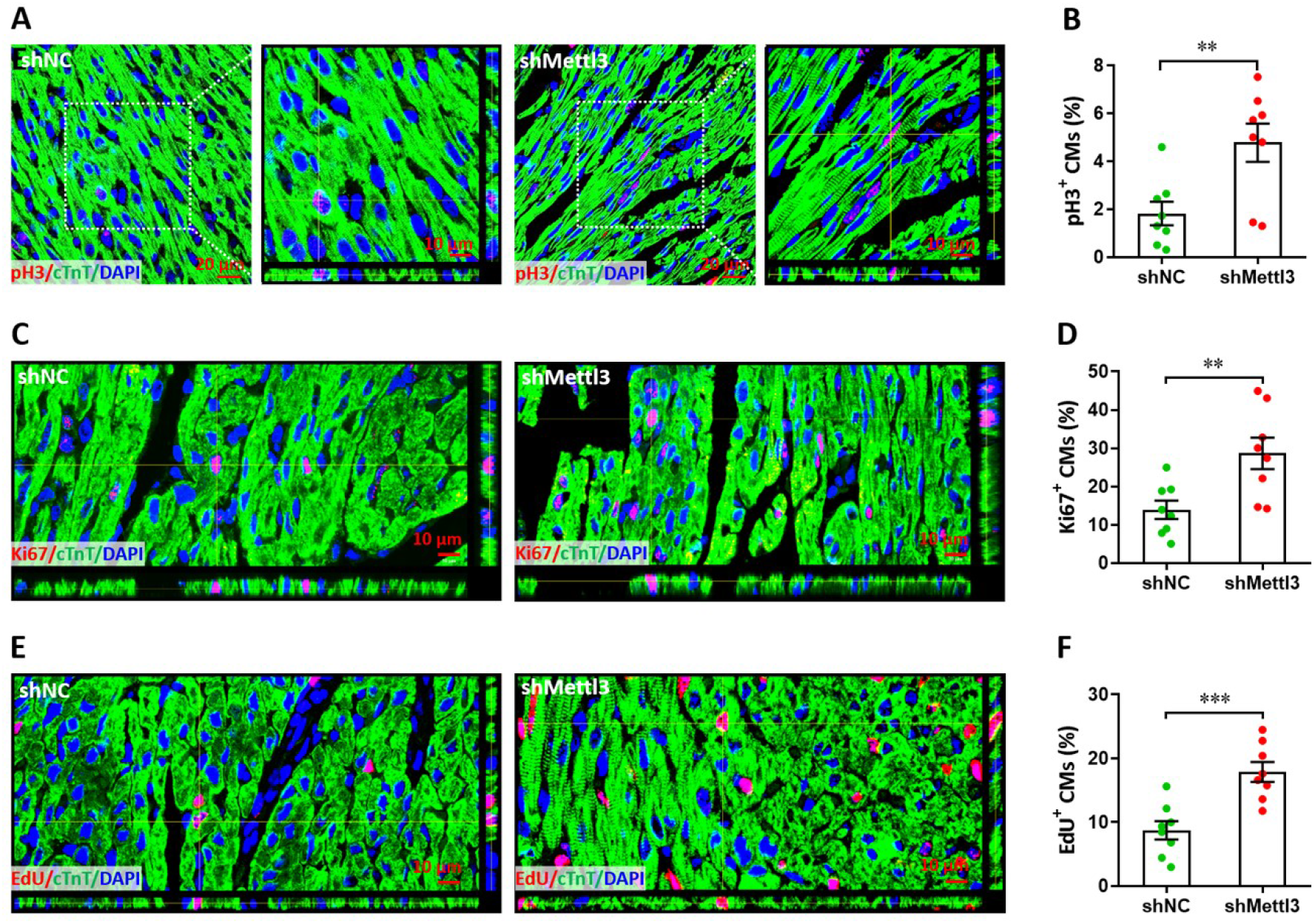
Mettl3 knockdown promotes cardiomyocyte proliferation in the remote area in injured heart at 5 dpr. (**A** and **B**) Representative confocal images (A) and quantification (B) of pH3^+^ cardiomyocytes in remote zone are shown (*n*=8 hearts per group). Right panel denotes the magnified Z-stack confocal images. (**C** and **D**) Representative images (C) and quantification (D) of Ki67^+^ cardiomyocytes in the remote zone are shown (*n*=8 hearts per group). (**E** and **F**) Representative Z-stack confocal images (E) and quantification (F) of EdU^+^ cardiomyocytes in remote zone are shown (*n*=8 hearts per group). All data are presented as the mean ± SEM, \*\**p*<0.01, \*\*\**p*<0.001 versus control. *P* values were determined by 2-tailed Student’s *t* test.

**Supplemental Figure 4.**
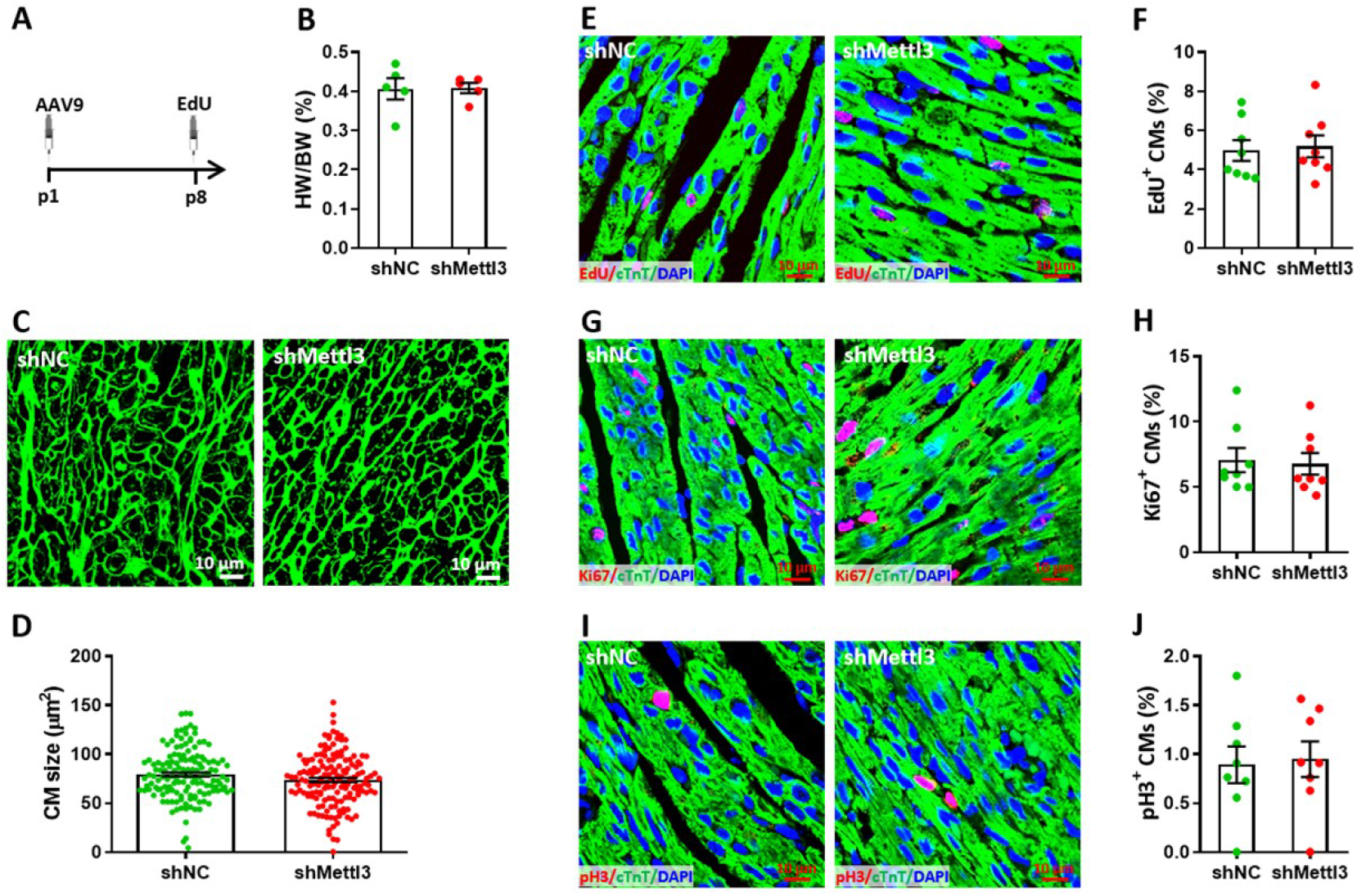
Effects of Mettl3 deficiency on cardiomyocyte proliferation in neonatal mice without injury at p8. (**A**) Schematic of AAV9-shMettl3 virus injection designed to knock down *Mettl3* in homeostatic neonatal hearts without injury. (**B**) Quantification of heart weight (HW) to body weight (BW) ratio (*n*=5 hearts per group). (**C**) Representative WGA staining images of myocardium in control and Mettl3-deficient hearts. (**D**) Quantification of cardiomyocyte size in control and Mettl3-deficient hearts are shown (*n*=∼150 cells from 5 hearts per group). (**E** and **F**) Representative images (E) and quantification (F) of EdU^+^ cardiomyocytes in ventricles at p8 (*n*=8 hearts per group). (**G** and **H**) Representative images (G) and quantification (H) of Ki67^+^ cardiomyocytes in ventricles at p8 (*n*=8 hearts per group). (**I** and **J**) Representative images (I) and quantification (J) of pH3^+^ cardiomyocytes in ventricles at p8 (*n*=8 hearts per group). All data are presented as the mean ± SEM.

**Supplemental Figure 5.**
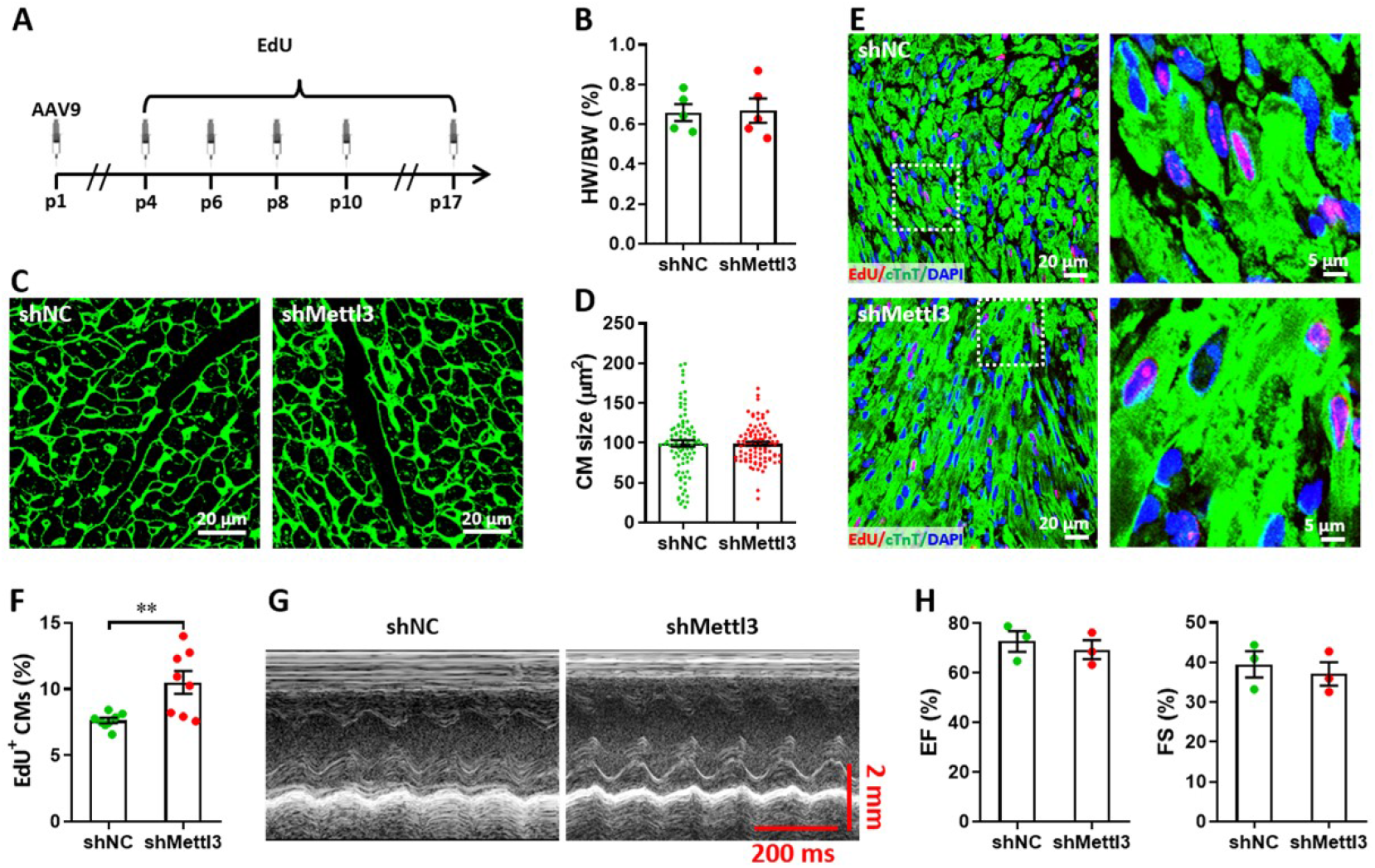
Effects of Mettl3 deficiency on heart regeneration in postnatal mice without injury at p17. (**A**) Schematic of AAV9-shMettl3 virus injection and EdU-pulse injection in homeostatic postnatal hearts without injury. (**B**) Quantification of heart weight (HW) to body weight (BW) ratio (*n*=5 hearts per group). (**C**) Representative WGA staining images of myocardium in control and Mettl3-deficient hearts. (**D**) Quantification of cardiomyocyte size in control and Mettl3-deficient hearts are shown (*n*=∼150 cells from 5 hearts per group). (**E** and **F**) Representative images (E) and quantification (F) of EdU^+^ cardiomyocytes in ventricles at p17 (*n*=8 hearts per group). (**G** and **H**) Representative images of M-model echocardiography (G) and quantification of LVEF (H, left) and LVFS (H, right) are shown (*n*=3 hearts per group). All data are presented as the mean ± SEM, \*\**p*<0.01 versus control. *P* values were determined by 2-tailed Student’s *t* test.

**Supplemental Figure 6.**
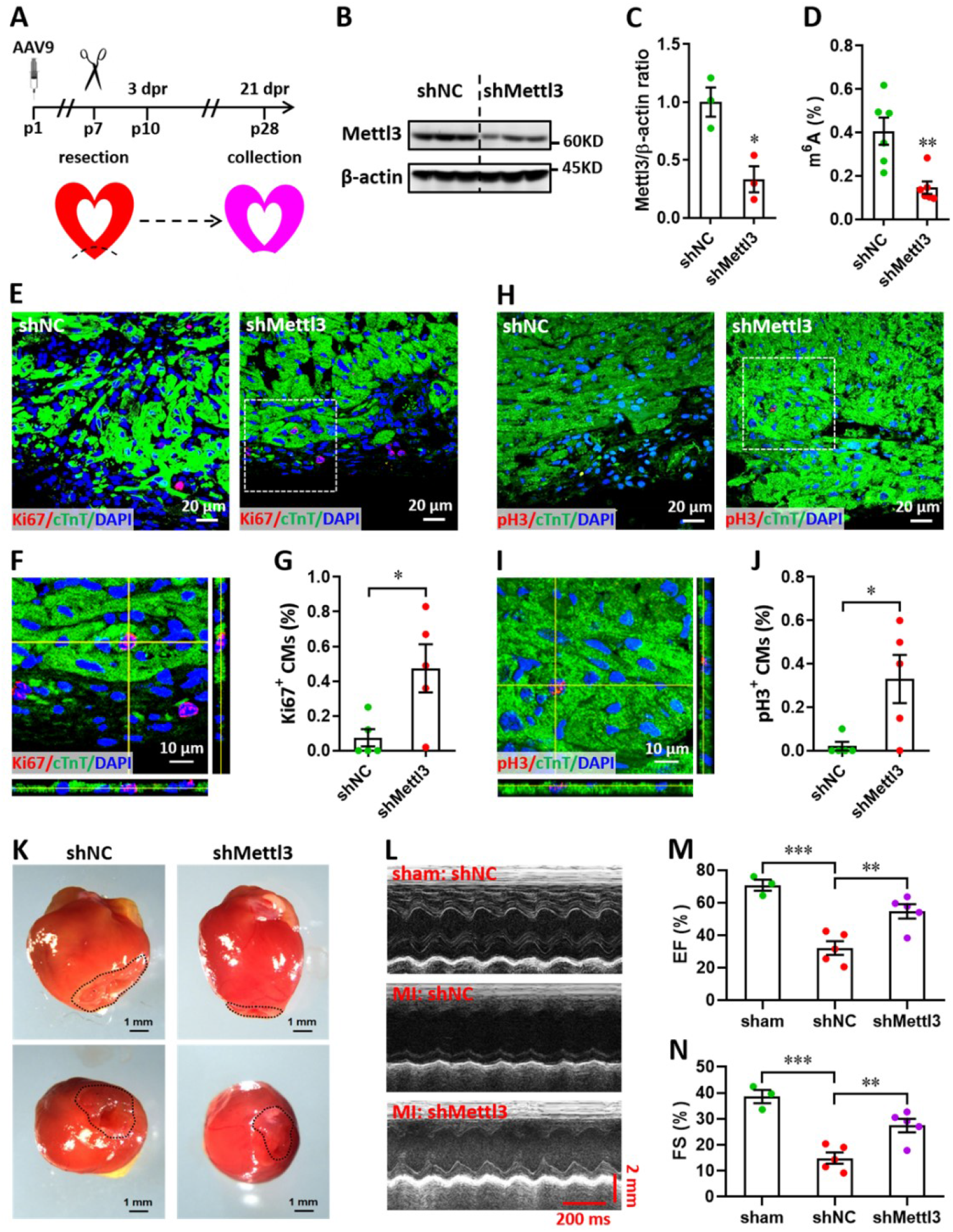
Mettl3 knockdown promotes heart regeneration upon apex resection at p7. (**A**) Schematic of AAV9-shMettl3 virus injection, apex resection, and sample collection. (**B** and **C**) Representative images (B) and quantification (C) of western blotting assay using heart tissues at p7 (*n*=3 hearts). (**D**) m^6^A levels in the injured heart at p7 were measured by ELISA-based quantification assay (*n*=6 hearts per group). (**E**-**G**) Representative images (E, F) and quantification (G) of Ki67^+^ cardiomyocytes in the border zone of apex are shown at 3 dpr (*n=*8 hearts). Representative Z-stack confocal image (F) denotes the area marked by dotted line in E panel. (**H**-**J**) Representative images (H, I) and quantification (J) of pH3^+^ cardiomyocytes in the border zone of apex are shown at 3 dpr (*n=*8 hearts). Representative Z-stack confocal image (I) denotes the area marked by dotted line in I panel. (**K**) Representative whole (upper panel) and apical (lower panel) images of neonatal hearts at 21 dpr. Dotted lines denote scars in cardiac apex. (**L**-**N**) Representative images of M-model echocardiography (L) and quantification of LVEF (M) and LVFS (N) at 21 dpr are shown (*n*=5 hearts). All data are presented as the mean ± SEM, \**p*<0.05, \*\**p*<0.01, \*\*\**p*<0.001. *P* values were determined by 2-tailed Student’s *t* test (C, D, G, and J), or by 1-way ANOVA with Dunnett’s multiple-comparison test (M and N).

**Supplemental Figure 7.**
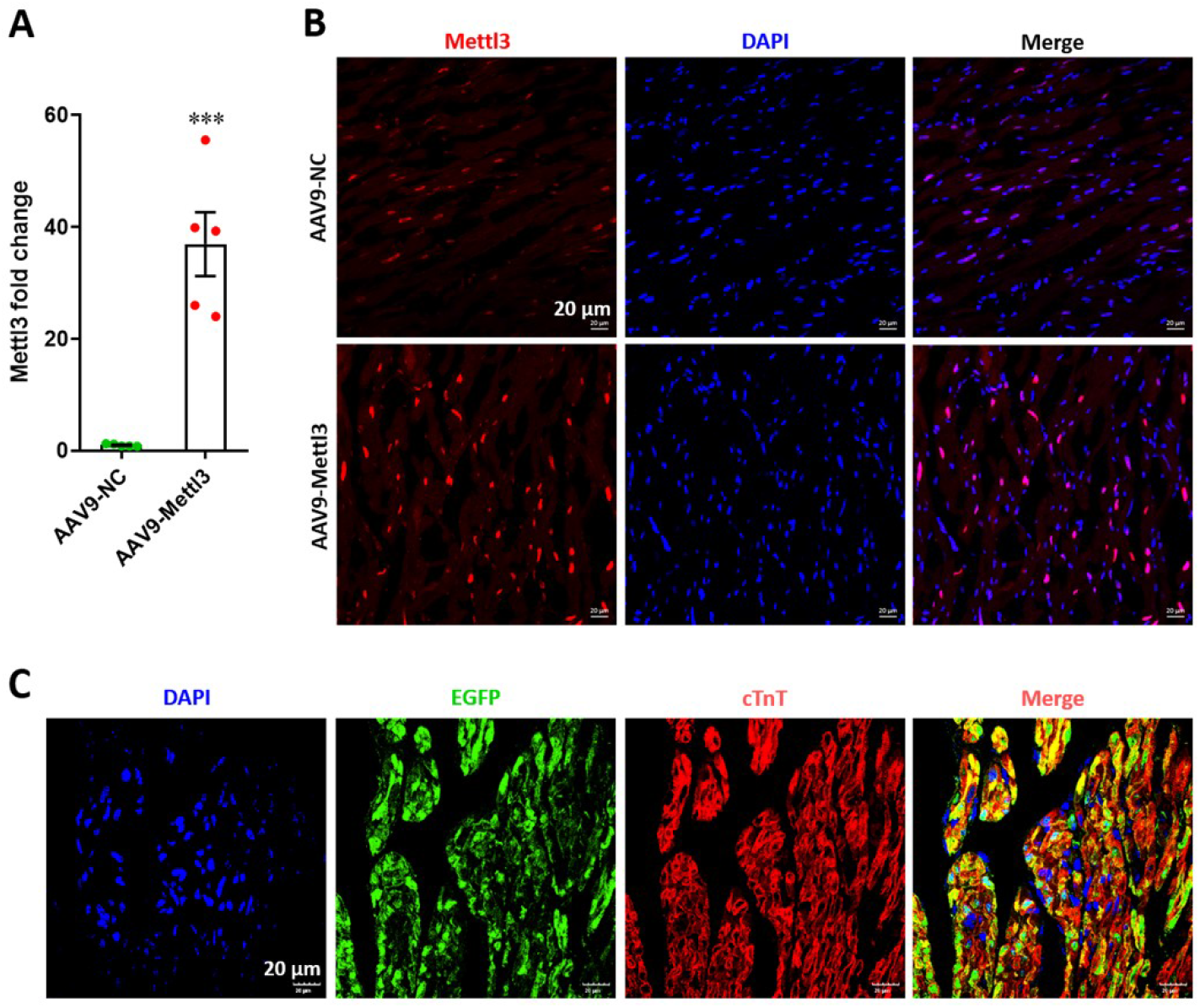
Overexpression of Mettl3 in postnatal hearts with AAV9 injection at p27. (**A**) qPCR validation of *Mettl3* mRNA expression in postnatal hearts at 27 dpr. Data are presented as the mean ± SEM (*n*=5 hearts per group), \*\*\**p*<0.001 versus control. *P* values were determined by 2-tailed Student’s *t* test. (**B**) Expression of Mettl3 protein was examined using immunofluorescence staining in postnatal hearts at 27 dpr. (**C**) Representative images of reporter gene (EGFP) expression mediated by AAV9-Mettl3 virus in heart at 27 dpr.

**Supplemental Figure 8.**
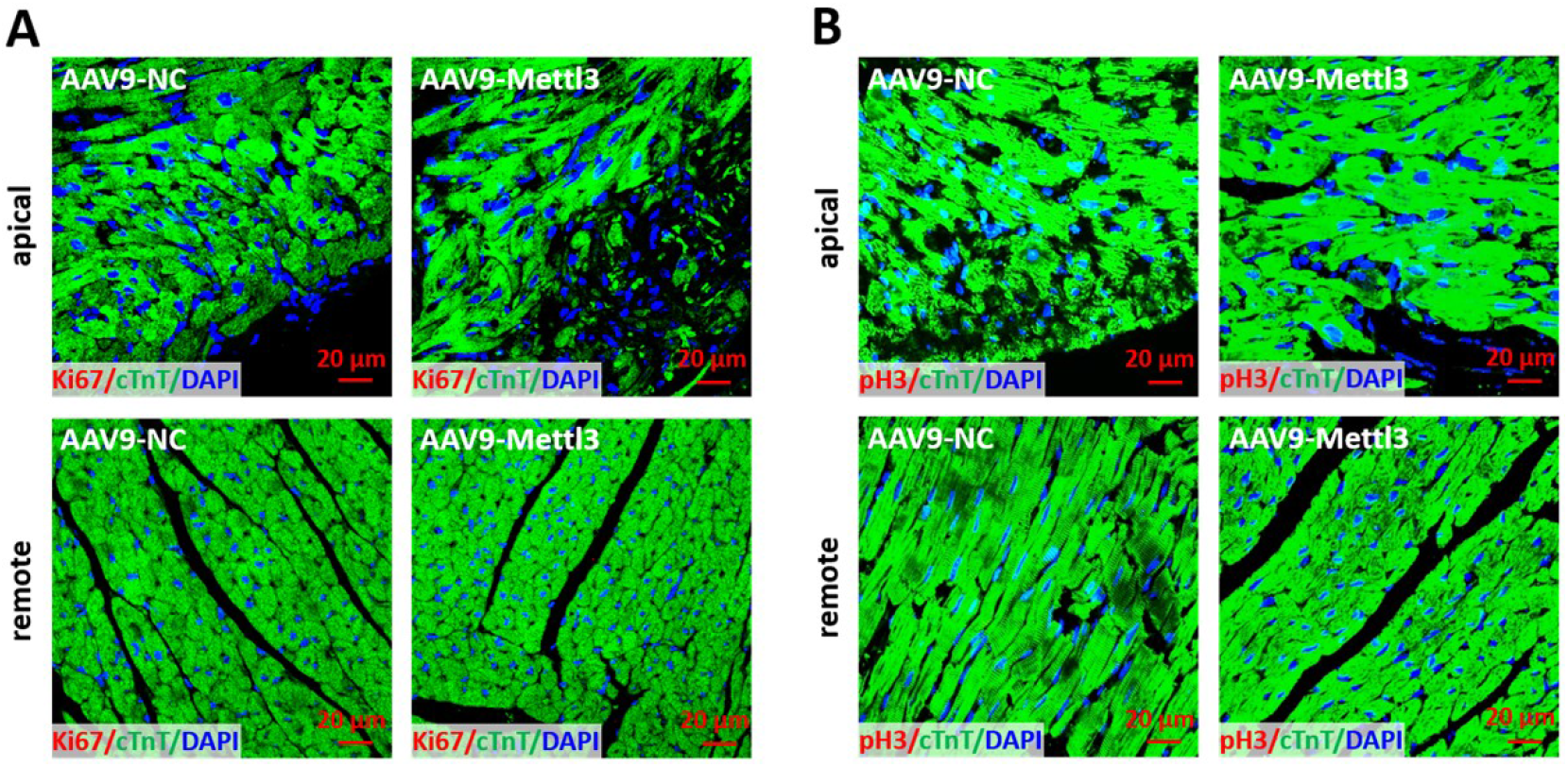
No proliferating cardiomyocytes was detected in heart at 27 dpr. (**A**) Sections from apical (upper panels) and remote (lower panels) ventricles were subjected to Ki67 and cTnT double staining. Representative images of cardiomyocytes in apical (upper panels) and remote (lower panels) zone showing that there was no Ki67^+^ cardiomyocytes in both groups at 27 dpr. (**B**) Sections from apical (upper panels) and remote (lower panels) ventricles were subjected to pH3 and cTnT double staining. Representative images of cardiomyocytes in apical (upper panels) and remote (lower panels) zone showing that there was no pH3^+^ cardiomyocytes in both groups at 27 dpr.

**Supplemental Figure 9.**
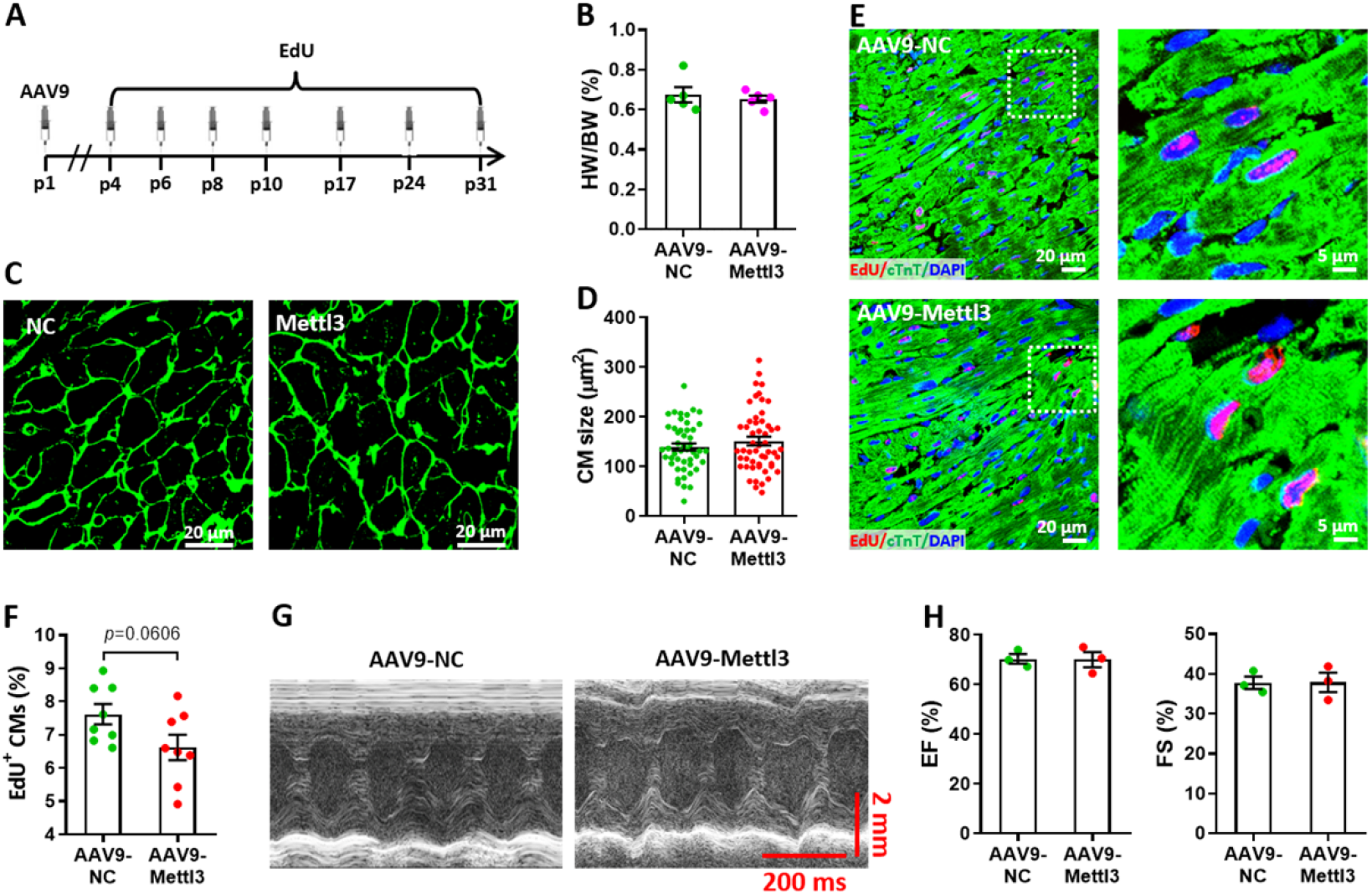
Effects of Mettl3 overexpression on heart regeneration in postnatal mice without injury at p31. (**A**) Schematic of AAV9-Mettl3 virus injection and EdU-pulse injection in homeostatic postnatal hearts without injury. (**B**) Quantification of heart weight (HW) to body weight (BW) ratio (*n*=5 hearts per group). (**C**) Representative WGA staining images of myocardium in control and Mettl3-deficient hearts. (**D**) Quantification of cardiomyocyte size in control and Mettl3-overexpressing hearts are shown (*n*=∼150 cells from 5 hearts per group). (**E** and **F**) Representative images (E) and quantification (F) of EdU^+^ cardiomyocytes in ventricles at p31 (*n*=8 hearts per group). (**G** and **H**) Representative images of M-model echocardiography (G) and quantification of LVEF (H, left) and LVFS (H, right) are shown (*n*=3 hearts per group). All data are presented as the mean ± SEM. *P* values were determined by 2-tailed Student’s *t* test.

**Supplemental Figure 10.**
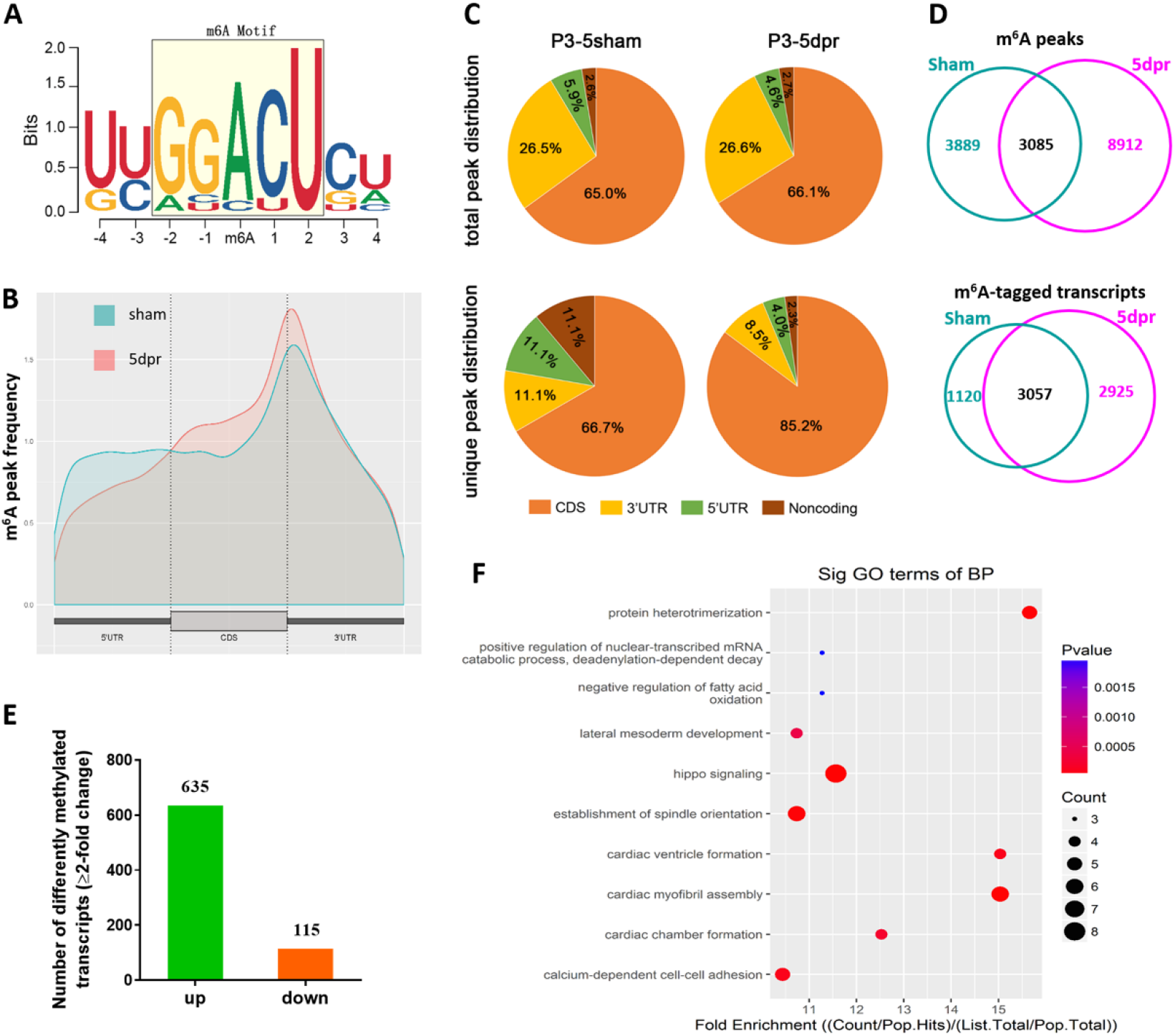
Variations of m^6^A-tagged transcripts in neonatal heart in response to injury. (**A**) Predominant consensus motif GGACU was detected basing on m^6^A-seq in both the sham and injured hearts at 5 dpr. (**B**) Distribution of m^6^A peaks along the 5’ untranslated region (5’UTR), coding region (CDS), and 3’ untranslated region (3’UTR) of total mRNAs from sham and apex-resected hearts. (**C**) Proportion of m^6^A peaks distribution in the 5’UTR, CDS, 3’UTR, and noncoding regions across the entire set of mRNA transcripts. Up panel denotes total peak distribution, down panel denotes unique peak distribution. (**D**) Venn diagrams showing the total m^6^A peaks (up panel) and m^6^A-tagged transcripts (low panel) between sham-operated and apex-resected hearts. (**E**) Numbers of the differently methylated genes with 2-fold or above change of m^6^A-tagging in the apex-resected hearts compared with sham. (**F**) A cluster profiler identified the enriched gene ontology processes of these 635 genes with upregulation of m^6^A modification.

**Supplemental Figure 11.**
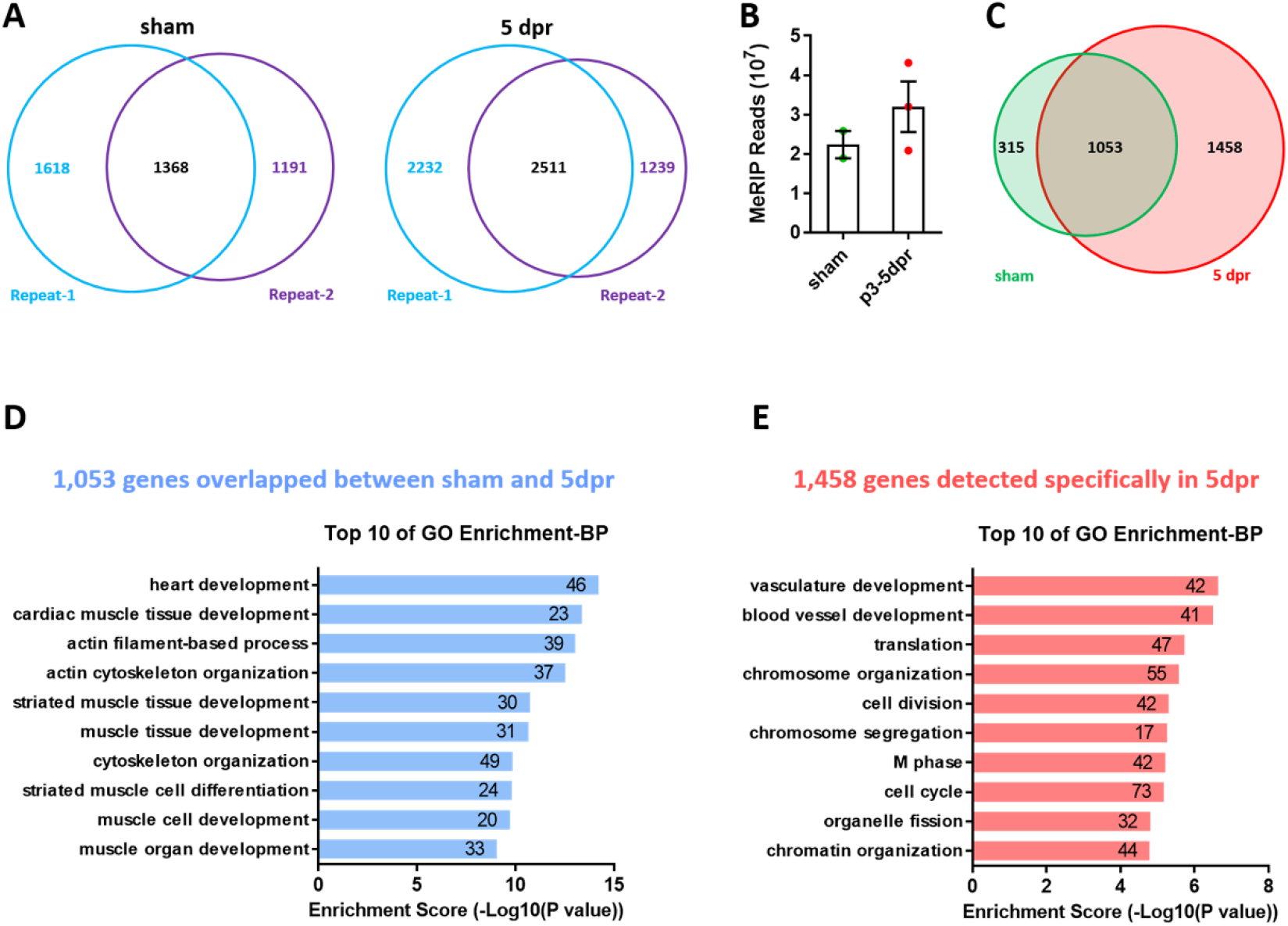
Apex resection upregulates levels of m^6^A-tagged transcripts in neonatal hearts. (**A**) Venn diagrams showing the overlap of m^6^A-tagged transcripts between two replicates in sham and 5dpr. (**B**) Total MeRIP bound reads from MeRIP-seq. (**C**) Venn diagrams of m^6^A-tagged transcripts identified by MeRIP-seq in neonatal hearts under sham and 5dpr conditions. (**D**) GO enrichment analyses of overlapped 1,053 genes between sham and 5dpr (related to C). (**E**) GO enrichment analyses of 1,458 genes specifically tagged in 5dpr (related to C).

**Supplemental Figure 12.**
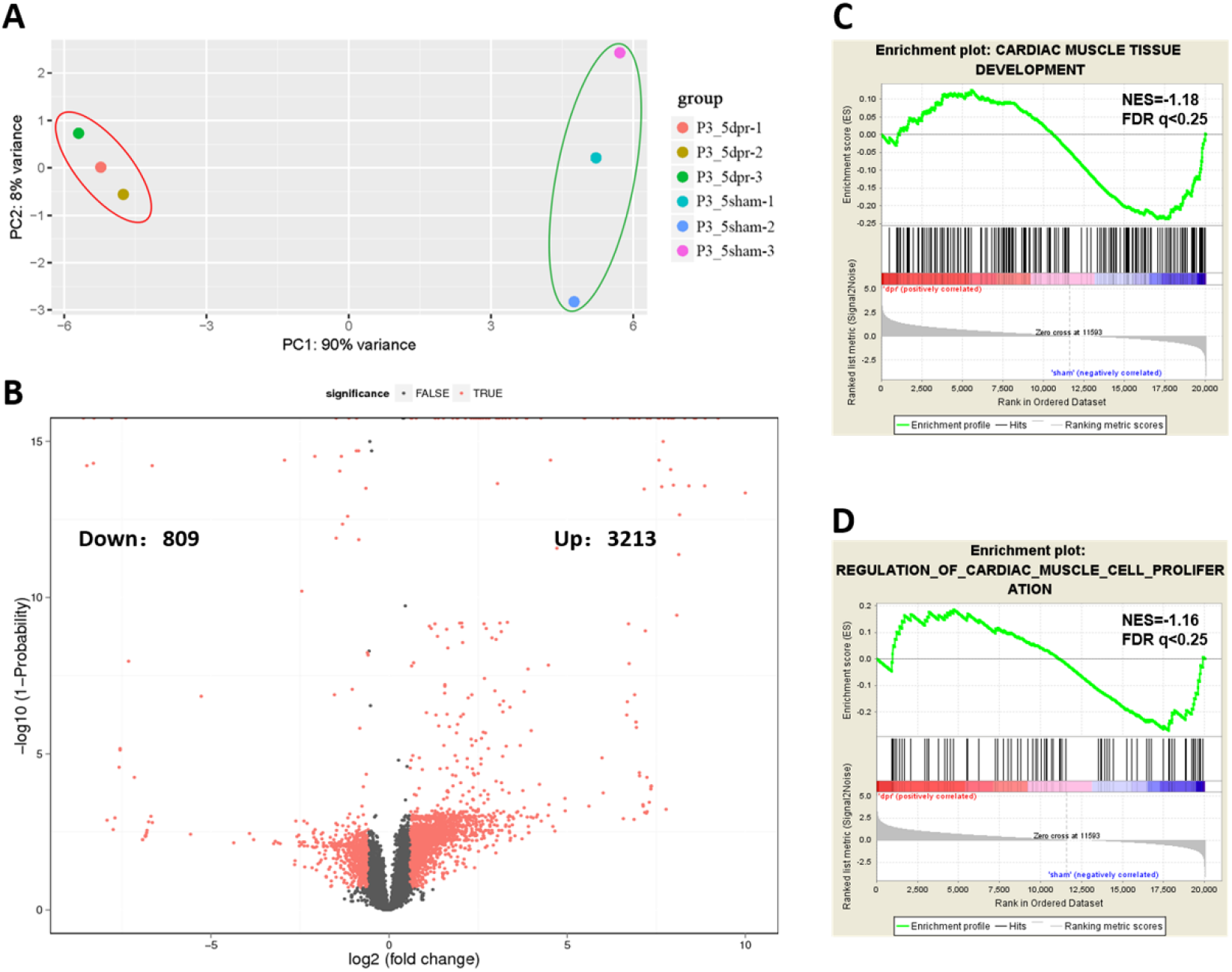
PCA, volcano plot, and GSEA analysis from RNA-seq data. (**A**) Principle component analysis (PCA) plot based on the RNA-seq data from control (sham) and injured hearts at 5 dpr. (**B**) Volcano plot for differentially expressed genes in the tests of neonatal hearts at 5 dpr versus uninjured controls. The red point in the plot represents the differentially expressed genes with statistical significance (FC > 1.5, Probability > 0.8). 3,213 upregulated and 809 downregulated genes passed the volcano plot filtering. (**C** and **D**) GSEA analysis based on RNA-seq reveals that cardiac muscle cell development (C) and proliferation (D) gene sets are related to neonatal heart regeneration.

**Supplemental Figure 13.**
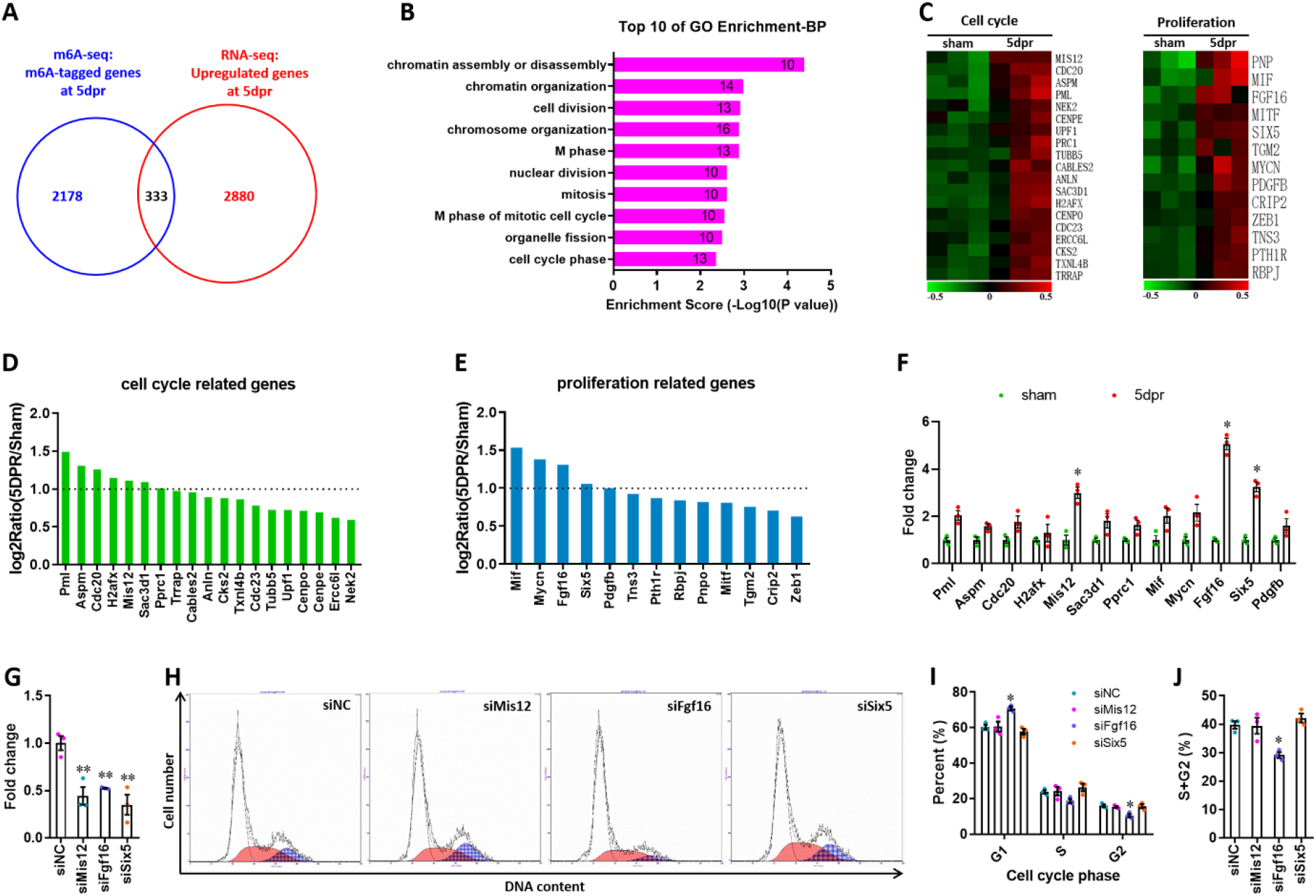
Identification of cell cycle and proliferation-related target genes from the overlapped gene sets between m^6^A-tagged and RNA-upregulated genes. (**A**) Venn diagrams showing the overlapped 333 genes between m^6^A-tagged and upregulated transcripts in neonatal hearts at 5dpr. (**B**) GO enrichment analyses of the overlapped 333 genes. (**C**) Heat map of mRNA expression of the cell cycle and proliferation-related genes from GO enrichment terms (*n*=3 hearts). (**D** and **E**) Quantification of mRNA expression of cell cycle- (D) and proliferation-related (E) genes from RNA-seq data. Dotted line indicates 2-fold upregulation. (**F**) Upregulated genes with more than 2-fold exchanges in RNA-seq (D and E) were further validated by qPCR in the injured hearts (*n*=5 hearts). (**G**) qPCR validation of siRNAs targeting for *Mis12*, *Fgf16*, and *Six5* genes in H9c2 cells (*n*=3). (**H**-**J**) H9c2 cells transfected with siRNAs for 48 hr were subjected to flow cytometry. Representative images (H) and quantification (I, J) of cell cycle distribution are shown (*n*=3). Data are presented as the mean ± SEM, \**p*<0.05, \*\**p*<0.01 versus sham (F) or controls (J, I, and J). *P* values were determined by 1-way (G and J) or 2-way (F and I) ANOVA with Dunnett’s multiple-comparison test.

**Supplemental Figure 14.**
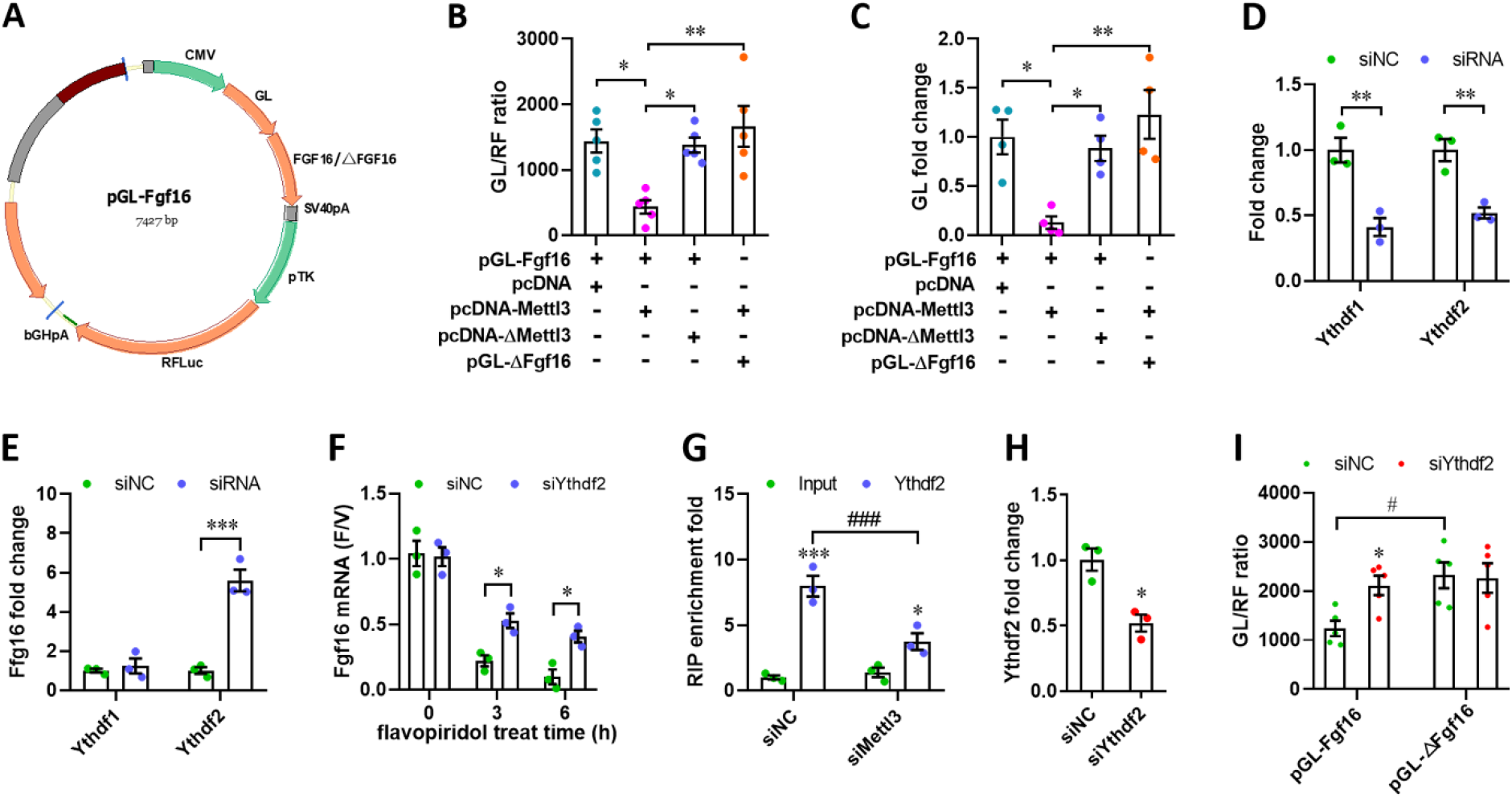
Mettl3 regulates Fgf16 expression in an Ythdf2-dependent manner. (**A**) Map of reporter plasmid for subcloning wild-type or m^6^A consensus sequence mutant Fgf16 cDNA to fuse with Gaussia luciferase reporter gene. (**B** and **C**) Quantification of relative luciferase activity (B, *n*=5) and mRNA level (C, *n*=4) in H9c2 cells with or without *Mettl3* overexpression. (**D**) qPCR validation of *Ythdf1* and *Ythdf2* silencing in primary cardiomyocytes (*n*=3). (**E**) qPCR validation of *Fgf16* in primary cardiomyocytes transfected with siYthdf1 or siYthdf2, respectively (*n*=3). (**F**) Primary cardiomyocytes were transfected with siRNAs for 48 hr and treated with vehicle or flavopiridol for 0 to 6 hr, followed by qPCR validation to determine the F/V ratio of *Fgf16* mRNA (*n*=3). (**G**) Ythdf2-RIP-qPCR analysis of *Fgf16* mRNA in primary cardiomyocytes transfected with siNC and siMettl3, respectively (*n*=3). (**H**) qPCR validation of *Ythdf2* silencing in H9c2 cells (*n*=3). (**I**) Wild-type and mutant reporter plasmids were respectively transfected into H9c2 cells treated with or without siYthdf2 for 48 hr. Relative luciferase activity was then quantified (*n*=5). All data are presented as the mean ± SEM, \**p*<0.05, \*\**p*<0.01, \*\*\**p*<0.001, ^#^*p*<0.05, ^###^*p*<0.001 versus control. *P* values were determined by 2-tailed Student’s *t* test (H), or by 1-way (B and C) or 2-way (D-G, and I) ANOVA with Dunnett’s multiple-comparison test.

## References

Arrell DK, Rosenow CS, Yamada S, Behfar A, Terzic A (2020). Cardiopoietic stem cell therapy restores infarction-altered cardiac proteome. NPJ Regen Med. 5, 5.

Barbieri I, Tzelepis K, Pandolfini L, Shi J, Millan-Zambrano G, Robson SC, Aspris D, Migliori V, Bannister AJ, Han N, De Braekeleer E, Ponstingl H, Hendrick A, Vakoc CR, Vassiliou GS, Kouzarides T (2017). Promoter-bound METTL3 maintains myeloid leukaemia by m(6)A-dependent translation control. Nature. 552, 126–131.

Batista PJ, Molinie B, Wang J, Qu K, Zhang J, Li L, Bouley DM, Lujan E, Haddad B, Daneshvar K, Carter AC, Flynn RA, Zhou C, Lim KS, Dedon P, Wernig M, Mullen AC, Xing Y, Giallourakis CC, Chang HY (2014). m(6)A RNA modification controls cell fate transition in mammalian embryonic stem cells. Cell Stem Cell. 15, 707–719.

Bergmann O, Bhardwaj RD, Bernard S, Zdunek S, Barnabe-Heider F, Walsh S, Zupicich J, Alkass K, Buchholz BA, Druid H, Jovinge S, Frisen J (2009). Evidence for Cardiomyocyte Renewal in Humans. Science. 324, 98–102.

Biressi S, Filareto A, Rando TA (2020). Stem cell therapy for muscular dystrophies. J Clin Invest. 130, 5652–5664.

Boccaletto P, Machnicka MA, Purta E, Piatkowski P, Baginski B, Wirecki TK, de Crecy-Lagard V, Ross R, Limbach PA, Kotter A, Helm M, Bujnicki JM (2018). MODOMICS: a database of RNA modification pathways. 2017 update. Nucleic Acids Res. 46, D303–D307.

Bokar JA, Shambaugh ME, Polayes D, Matera AG, Rottman FM (1997). Purification and cDNA cloning of the AdoMet-binding subunit of the human mRNA (N6-adenosine)-methyltransferase. RNA. 3, 1233–1247.

Brundel BJ, Van Gelder IC, Henning RH, Tuinenburg AE, Wietses M, Grandjean JG, Wilde AA, Van Gilst WH, Crijns HJ (2001). Alterations in potassium channel gene expression in atria of patients with persistent and paroxysmal atrial fibrillation: differential regulation of protein and mRNA levels for K+ channels. J Am Coll Cardiol. 37, 926–932.

Bui AL, Horwich TB, Fonarow GC (2011). Epidemiology and risk profile of heart failure. Nat Rev Cardiol. 8, 30–41.

Chen M, Wei L, Law CT, Tsang FH, Shen J, Cheng CL, Tsang LH, Ho DW, Chiu DK, Lee JM, Wong CC, Ng IO, Wong CM (2018). RNA N6-methyladenosine methyltransferase-like 3 promotes liver cancer progression through YTHDF2-dependent posttranscriptional silencing of SOCS2. Hepatology. 67, 2254–2270.

Chong JJ, Yang X, Don CW, Minami E, Liu YW, Weyers JJ, Mahoney WM, Van Biber B, Cook SM, Palpant NJ, Gantz JA, Fugate JA, Muskheli V, Gough GM, Vogel KW, Astley CA, Hotchkiss CE, Baldessari A, Pabon L, Reinecke H, Gill EA, Nelson V, Kiem HP, Laflamme MA, Murry CE (2014). Human embryonic-stem-cell-derived cardiomyocytes regenerate non-human primate hearts. Nature. 510, 273–277.

Dominissini D, Moshitch-Moshkovitz S, Schwartz S, Salmon-Divon M, Ungar L, Osenberg S, Cesarkas K, Jacob-Hirsch J, Amariglio N, Kupiec M, Sorek R, Rechavi G (2012). Topology of the human and mouse m6A RNA methylomes revealed by m6A-seq. Nature. 485, 201–206.

Dorn LE, Lasman L, Chen J, Xu X, Hund TJ, Medvedovic M, Hanna JH, van Berlo JH, Accornero F (2019). The N(6)-Methyladenosine mRNA Methylase METTL3 Controls Cardiac Homeostasis and Hypertrophy. Circulation. 139, 533–545.

Du H, Zhao Y, He J, Zhang Y, Xi H, Liu M, Ma J, Wu L (2016). YTHDF2 destabilizes m(6)A-containing RNA through direct recruitment of the CCR4-NOT deadenylase complex. Nat Commun. 7, 12626.

Eisen MB, Spellman PT, Brown PO, Botstein D (1998). Cluster analysis and display of genome-wide expression patterns. Proc Natl Acad Sci U S A. 95, 14863–14868.

Feng T, Meng J, Kou S, Jiang Z, Huang X, Lu Z, Zhao H, Lau LF, Zhou B, Zhang H (2019). CCN1-Induced Cellular Senescence Promotes Heart Regeneration. Circulation. 139, 2495–2498.

Frye M, Harada BT, Behm M, He C (2018). RNA modifications modulate gene expression during development. Science. 361, 1346–1349.

Fu Y, Dominissini D, Rechavi G, He C (2014). Gene expression regulation mediated through reversible m(6)A RNA methylation. Nat Rev Genet. 15, 293–306.

Geula S, Moshitch-Moshkovitz S, Dominissini D, Mansour AA, Kol N, Salmon-Divon M, Hershkovitz V, Peer E, Mor N, Manor YS, Ben-Haim MS, Eyal E, Yunger S, Pinto Y, Jaitin DA, Viukov S, Rais Y, Krupalnik V, Chomsky E, Zerbib M, Maza I, Rechavi Y, Massarwa R, Hanna S, Amit I, Levanon EY, Amariglio N, Stern-Ginossar N, Novershtern N, Rechavi G, Hanna JH (2015). Stem cells. m6A mRNA methylation facilitates resolution of naive pluripotency toward differentiation. Science. 347, 1002–1006.

Haussmann IU, Bodi Z, Sanchez-Moran E, Mongan NP, Archer N, Fray RG, Soller M (2016). m(6)A potentiates Sxl alternative pre-mRNA splicing for robust Drosophila sex determination. Nature. 540, 301–304.

Hotta Y, Sasaki S, Konishi M, Kinoshita H, Kuwahara K, Nakao K, Itoh N (2008). Fgf16 Is Required for Cardiomyocyte Proliferation in the Mouse Embryonic Heart. Dev Dynam. 237, 2947–2954.

Hsu PJ, Shi H, He C (2017). Epitranscriptomic influences on development and disease. Genome Biol. 18, 197.

Ivanova I, Much C, Di Giacomo M, Azzi C, Morgan M, Moreira PN, Monahan J, Carrieri C, Enright AJ, O’Carroll D (2017). The RNA m(6)A Reader YTHDF2 Is Essential for the Post-transcriptional Regulation of the Maternal Transcriptome and Oocyte Competence. Mol Cell. 67, 1059–1067 e1054.

Kim D, Landmead B, Salzberg SL (2015). HISAT: a fast spliced aligner with low memory requirements. Nat Methods. 12, 357–U121.

Lazar E, Sadek HA, Bergmann O (2017). Cardiomyocyte renewal in the human heart: insights from the fall-out. Eur Heart J. 38, 2333–2342.

Li B, Dewey CN (2011). RSEM: accurate transcript quantification from RNA-Seq data with or without a reference genome. BMC Bioinformatics. 12, 323.

Li F, Yi Y, Miao Y, Long W, Long T, Chen S, Cheng W, Zou C, Zheng Y, Wu X, Ding J, Zhu K, Chen D, Xu Q, Wang J, Liu Q, Zhi F, Ren J, Cao Q, Zhao W (2019). N(6)-Methyladenosine Modulates Nonsense-Mediated mRNA Decay in Human Glioblastoma. Cancer Res. 79, 5785–5798.

Lin Z, Pu WT (2014). Strategies for cardiac regeneration and repair. Sci Transl Med. 6, 239rv231.

Litvinukova M, Talavera-Lopez C, Maatz H, Reichart D, Worth CL, Lindberg EL, Kanda M, Polanski K, Heinig M, Lee M, Nadelmann ER, Roberts K, Tuck L, Fasouli ES, DeLaughter DM, McDonough B, Wakimoto H, Gorham JM, Samari S, Mahbubani KT, Saeb-Parsy K, Patone G, Boyle JJ, Zhang H, Viveiros A, Oudit GY, Bayraktar OA, Seidman JG, Seidman CE, Noseda M, Hubner N, Teichmann SA (2020). Cells of the adult human heart. Nature. 588, 466–472.

Liu J, Yue Y, Han D, Wang X, Fu Y, Zhang L, Jia G, Yu M, Lu Z, Deng X, Dai Q, Chen W, He C (2014). A METTL3-METTL14 complex mediates mammalian nuclear RNA N6-adenosine methylation. Nat Chem Biol. 10, 93–95.

Lu SY, Sheikh F, Sheppard PC, Fresnoza A, Duckworth ML, Detillieux KA, Cattini PA (2008). FGF-16 is required for embryonic heart development. Biochem Bioph Res Co. 373, 270–274.

Luo GZ, MacQueen A, Zheng G, Duan H, Dore LC, Lu Z, Liu J, Chen K, Jia G, Bergelson J, He C (2014). Unique features of the m6A methylome in Arabidopsis thaliana. Nat Commun. 5, 5630.

Mahmoud AI, Kocabas F, Muralidhar SA, Kimura W, Koura AS, Thet S, Porrello ER, Sadek HA (2013). Meis1 regulates postnatal cardiomyocyte cell cycle arrest. Nature. 497, 249–253.

Mathiyalagan P, Adamiak M, Mayourian J, Sassi Y, Liang Y, Agarwal N, Jha D, Zhang S, Kohlbrenner E, Chepurko E, Chen J, Trivieri MG, Singh R, Bouchareb R, Fish K, Ishikawa K, Lebeche D, Hajjar RJ, Sahoo S (2019). FTO-Dependent N(6)-Methyladenosine Regulates Cardiac Function During Remodeling and Repair. Circulation. 139, 518–532.

Meng J, Lu Z, Liu H, Zhang L, Zhang S, Chen Y, Rao MK, Huang Y (2014). A protocol for RNA methylation differential analysis with MeRIP-Seq data and exomePeak R/Bioconductor package. Methods. 69, 274–281.

Meyer KD, Saletore Y, Zumbo P, Elemento O, Mason CE, Jaffrey SR (2012). Comprehensive Analysis of mRNA Methylation Reveals Enrichment in 3’ UTRs and near Stop Codons. Cell. 149, 1635–1646.

Nakada Y, Canseco DC, Thet S, Abdisalaam S, Asaithamby A, Santos CX, Shah AM, Zhang H, Faber JE, Kinter MT, Szweda LI, Xing C, Hu Z, Deberardinis RJ, Schiattarella G, Hill JA, Oz O, Lu Z, Zhang CC, Kimura W, Sadek HA (2017). Hypoxia induces heart regeneration in adult mice. Nature. 541, 222–227.

Narula J, Haider N, Virmani R, DiSalvo TG, Kolodgie FD, Hajjar RJ, Schmidt U, Semigran MJ, Dec GW, Khaw BA (1996). Apoptosis in myocytes in end-stage heart failure. N Engl J Med. 335, 1182–1189.

Paris J, Morgan M, Campos J, Spencer GJ, Shmakova A, Ivanova I, Mapperley C, Lawson H, Wotherspoon DA, Sepulveda C, Vukovic M, Allen L, Sarapuu A, Tavosanis A, Guitart AV, Villacreces A, Much C, Choe J, Azar A, van de Lagemaat LN, Vernimmen D, Nehme A, Mazurier F, Somervaille TCP, Gregory RI, O’Carroll D, Kranc KR (2019). Targeting the RNA m(6)A Reader YTHDF2 Selectively Compromises Cancer Stem Cells in Acute Myeloid Leukemia. Cell Stem Cell.

Percie du Sert N, Hurst V, Ahluwalia A, Alam S, Avey MT, Baker M, Browne WJ, Clark A, Cuthill IC, Dirnagl U, Emerson M, Garner P, Holgate ST, Howells DW, Karp NA, Lazic SE, Lidster K, MacCallum CJ, Macleod M, Pearl EJ, Petersen OH, Rawle F, Reynolds P, Rooney K, Sena ES, Silberberg SD, Steckler T, Wurbel H (2020). The ARRIVE guidelines 2.0: Updated guidelines for reporting animal research. PLoS Biol. 18, e3000410.

Porrello ER, Mahmoud AI, Simpson E, Hill JA, Richardson JA, Olson EN, Sadek HA (2011). Transient Regenerative Potential of the Neonatal Mouse Heart. Science. 331, 1078–1080.

Porrello ER, Mahmoud AI, Simpson E, Johnson BA, Grinsfelder D, Canseco D, Mammen PP, Rothermel BA, Olson EN, Sadek HA (2013). Regulation of neonatal and adult mammalian heart regeneration by the miR-15 family. P Natl Acad Sci USA. 110, 187–192.

Preissl S, Schwaderer M, Raulf A, Hesse M, Gruning BA, Kobele C, Backofen R, Fleischmann BK, Hein L, Gilsbach R (2015). Deciphering the Epigenetic Code of Cardiac Myocyte Transcription. Circ Res. 117, 413–423.

Qi XF, Chen ZY, Xia JB, Zheng L, Zhao H, Pi LQ, Park KS, Kim SK, Lee KJ, Cai DQ (2015). FoxO3a suppresses the senescence of cardiac microvascular endothelial cells by regulating the ROS-mediated cell cycle. J Mol Cell Cardiol. 81, 114–126.

Roundtree IA, Evans ME, Pan T, He C (2017). Dynamic RNA Modifications in Gene Expression Regulation. Cell. 169, 1187–1200.

Sahara M, Santoro F, Chien KR (2015). Programming and reprogramming a human heart cell. EMBO J. 34, 710–738.

Sander V, Sune G, Jopling C, Morera C, Izpisua Belmonte JC (2013). Isolation and in vitro culture of primary cardiomyocytes from adult zebrafish hearts. Nat Protoc. 8, 800–809.

Scott L, Fender AC, Saljic A, Li LG, Chen XH, Wang XL, Linz D, Lang JL, Hohl M, Twomey D, Pham TT, Diaz-Lankenau R, Chelu MG, Kamler M, Entman ML, Taffet GE, Sanders P, Dobrev D, Li N (2021). NLRP3 inflammasome is a key driver of obesity-induced atrial arrhythmias. Cardiovascular Research. 117, 1746–1759.

Shi H, Wei J, He C (2019). Where, When, and How: Context-Dependent Functions of RNA Methylation Writers, Readers, and Erasers. Mol Cell. 74, 640–650.

Shi H, Zhang X, Weng YL, Lu Z, Liu Y, Li J, Hao P, Zhang Y, Zhang F, Wu Y, Delgado JY, Su Y, Patel MJ, Cao X, Shen B, Huang X, Ming GL, Zhuang X, Song H, He C, Zhou T (2018). m(6)A facilitates hippocampus-dependent learning and memory through YTHDF1. Nature. 563, 249–253.

Su YR, Chiusa M, Brittain E, Hemnes AR, Absi TS, Lim CC, Di Salvo TG (2015). Right ventricular protein expression profile in end-stage heart failure. Pulm Circ. 5, 481–497.

Tarazona S, Garcia-Alcalde F, Dopazo J, Ferrer A, Conesa A (2011). Differential expression in RNA-seq: a matter of depth. Genome Res. 21, 2213–2223.

Vu LP, Pickering BF, Cheng Y, Zaccara S, Nguyen D, Minuesa G, Chou T, Chow A, Saletore Y, MacKay M, Schulman J, Famulare C, Patel M, Klimek VM, Garrett-Bakelman FE, Melnick A, Carroll M, Mason CE, Jaffrey SR, Kharas MG (2017). The N(6)-methyladenosine (m(6)A)-forming enzyme METTL3 controls myeloid differentiation of normal hematopoietic and leukemia cells. Nat Med. 23, 1369–1376.

Wang CX, Cui GS, Liu X, Xu K, Wang M, Zhang XX, Jiang LY, Li A, Yang Y, Lai WY, Sun BF, Jiang GB, Wang HL, Tong WM, Li W, Wang XJ, Yang YG, Zhou Q (2018a). METTL3-mediated m6A modification is required for cerebellar development. PLoS Biol. 16, e2004880.

Wang H, Zuo H, Liu J, Wen F, Gao Y, Zhu X, Liu B, Xiao F, Wang W, Huang G, Shen B, Ju Z (2018b). Loss of YTHDF2-mediated m(6)A-dependent mRNA clearance facilitates hematopoietic stem cell regeneration. Cell Res. 28, 1035–1038.

Wang X, Lu Z, Gomez A, Hon GC, Yue Y, Han D, Fu Y, Parisien M, Dai Q, Jia G, Ren B, Pan T, He C (2014a). N6-methyladenosine-dependent regulation of messenger RNA stability. Nature. 505, 117–120.

Wang X, Zhao BS, Roundtree IA, Lu Z, Han D, Ma H, Weng X, Chen K, Shi H, He C (2015). N(6)-methyladenosine Modulates Messenger RNA Translation Efficiency. Cell. 161, 1388–1399.

Wang Y, Li Y, Toth JI, Petroski MD, Zhang Z, Zhao JC (2014b). N6-methyladenosine modification destabilizes developmental regulators in embryonic stem cells. Nat Cell Biol. 16, 191–198.

Wei LH, Song P, Wang Y, Lu Z, Tang Q, Yu Q, Xiao Y, Zhang X, Duan HC, Jia G (2018). The m(6)A Reader ECT2 Controls Trichome Morphology by Affecting mRNA Stability in Arabidopsis. Plant Cell. 30, 968–985.

Weng YL, Wang X, An R, Cassin J, Vissers C, Liu YY, Liu YJ, Xu TL, Wang XY, Wong SZH, Joseph J, Dore LC, Dong Q, Zheng W, Jin P, Wu H, Shen B, Zhuang XX, He C, Liu K, Song HJ, Ming GL (2018). Epitranscriptomic m(6)A Regulation of Axon Regeneration in the Adult Mammalian Nervous System. Neuron. 97, 313-+.

Wu HY, Zhou YM, Liao ZQ, Zhong JW, Liu YB, Zhao H, Liang CQ, Huang RJ, Park KS, Feng SS, Zheng L, Cai DQ, Qi XF (2021). Fosl1 is vital to heart regeneration upon apex resection in adult Xenopus tropicalis. NPJ Regen Med. 6, 36.

Xin B, Tao F, Wang Y, Liu H, Ma C, Xu P (2017). Coordination of metabolic pathways: Enhanced carbon conservation in 1,3-propanediol production by coupling with optically pure lactate biosynthesis. Metab Eng. 41, 102–114.

Yu W, Huang X, Tian X, Zhang H, He L, Wang Y, Nie Y, Hu S, Lin Z, Zhou B, Pu W, Lui KO (2016). GATA4 regulates Fgf16 to promote heart repair after injury. Development. 143, 936–949.

Zhao BS, Wang X, Beadell AV, Lu Z, Shi H, Kuuspalu A, Ho RK, He C (2017). m(6)A-dependent maternal mRNA clearance facilitates zebrafish maternal-to-zygotic transition. Nature. 542, 475–478.

Zhou DC, Su YH, Jiang FQ, Xia JB, Wu HY, Chang ZS, Peng WT, Song GH, Park KS, Kim SK, Cai DQ, Zheng L, Qi XF (2018). CpG oligodeoxynucleotide preconditioning improves cardiac function after myocardial infarction via modulation of energy metabolism and angiogenesis. J Cell Physiol. 233, 4245–4257.

